# Assembly defects of the human tRNA splicing endonuclease contribute to impaired pre-tRNA processing in pontocerebellar hypoplasia

**DOI:** 10.1101/2020.08.03.234229

**Authors:** Samoil Sekulovski, Pascal Devant, Silvia Panizza, Tasos Gogakos, Anda Pitiriciu, Katharina Heitmeier, Ewan Phillip Ramsay, Marie Barth, Carla Schmidt, Stefan Weitzer, Thomas Tuschl, Frank Baas, Javier Martinez, Simon Trowitzsch

## Abstract

Introns of human transfer RNA precursors (pre-tRNAs) are excised by the tRNA splicing endonuclease TSEN in complex with the RNA kinase CLP1. Mutations in TSEN/CLP1 occur in patients with pontocerebellar hypoplasia (PCH), however, their role in the disease is unclear. Here, we show that intron excision is catalyzed by tetrameric TSEN assembled from inactive heterodimers independently of CLP1. Splice site recognition involves the mature domain and the anticodon-intron base pair of pre-tRNAs. The 2.1-Å resolution X-ray crystal structure of a TSEN15–34 heterodimer and differential scanning fluorimetry analyses show that PCH mutations cause thermal destabilization. While endonuclease activity in recombinant mutant TSEN is unaltered, we observe assembly defects and reduced pre-tRNA cleavage activity resulting in an imbalanced pre-tRNA pool in PCH patient-derived fibroblasts. Our work defines the molecular principles of intron excision in humans and provides evidence that modulation of TSEN stability may contribute to PCH phenotypes.

## Main text

All nuclear-encoded transfer RNAs (tRNAs) are processed and modified during trafficking to the cytoplasm, to create functional, aminoacylated tRNAs ^1^. In humans, 28 out of 429 predicted high confidence tRNA genes contain introns that must be removed from precursor tRNAs (pre-tRNAs) by splicing ^2,3^ (http://gtrnadb.ucsc.edu/). Some isodecoders, e.g. tRNA^Tyr^_GTA_, tRNA^Ile^_TAT_, and tRNA^Leu^_CAA_, are only encoded as intron-containing precursors, for which splicing is essential for their production ^4^. Intron excision and ligation of the 5’ and 3’ tRNA exons is catalyzed by two multiprotein assemblies: the tRNA splicing endonuclease (TSEN) ^5^ and the tRNA ligase complex ^6^, respectively.

The human TSEN complex consists of two catalytic subunits, TSEN2 and TSEN34, and two structural subunits, TSEN15 and TSEN54, all expressed at very low copy numbers of ~ 100 molecules per cell ^5,7^. TSEN2–54 and TSEN15–34 are inferred to form distinct heterodimers from yeast-two-hybrid experiments, however a solution NMR structure has challenged the proposed model of TSEN assembly ^8,9^. Based on their quaternary structure, archaeal tRNA endonucleases have been classified into four types, α_4_, α’_2_, (αβ)_2_, and ε_2_ ^10,11^, whereas the eukaryotic tRNA endonucleases adapt a heterotetrameric αβγδ arrangement ^5,8^. Homotetramer formation in archaeal α_4_-type endonucleases is mediated by a hydrophobic domain interface involving antiparallel β strands of two neighboring α subunits and interactions between a negatively charged L10 loop of one α subunit with a positively charged pocket of an opposing α subunit. These interactions are conserved in all four types of archaeal endonucleases and were also suggested to occur in eukaryotic endonucleases. In humans, TSEN2 and TSEN34 are each predicted to harbor a catalytic triad, composed of Tyr^369^/His^377^/Lys^416^ in TSEN2 and Tyr^247^/His^255^/Lys^286^ in TSEN34, responsible for cleavage at the 5’ and 3’ splice sites, respectively ^5,8,12^. His^377^ and His^255^ are supposed to act as general acids at the scissile phosphates of the exon-intron junctions of pre-tRNAs ^5,12,13^. Furthermore, TSEN54 was suggested to function as a ‘molecular ruler’ measuring the distance from the mature domain of the tRNA to define the 5’ splice site ^8,13–16^.

The intron in pre-tRNAs is suggested to allow the formation of a double helix that extends the anticodon stem in the conventional tRNA cloverleaf structure and presents the 5’ and 3’ splice sites in single-stranded regions accessible for cleavage ^17,18^. Such a bulge-helix-bulge (BHB) conformation was postulated to act as a universal recognition motif in archaeal pre-tRNA splicing allowing for intron recognition at various positions in pre-tRNAs ^19^. In contrast, eukaryotic introns strictly locate one nucleotide 3’ to the anticodon triplet in the anticodon loop with varying sequence and length ^3,14^. Experiments using yeast and *Xenopus* tRNA endonucleases showed that cleavage at the exon-intron boundaries requires the presence of an anticodon–intron (A–I) base pair that controls cleavage at the 3’ splice site besides positioning of the 5’ splice site via the mature domain of the pre-tRNA ^14,20,21^. The X-ray crystal structure of an archaeal endonuclease with a BHB-substrate showed that two arginine residues at each active site form a cation-π sandwich with a flipped-out purine base of the pre-tRNA and thereby fixing the substrate for an S_N_2-type in line-attack ^12,13,16^. However, biochemical experiments using the yeast endonuclease showed that the cation-π sandwich is only required for cleavage at the 5’ splice site, whereas it is dispensable for catalysis at the 3’ splice site ^12^.

Specific to mammals is the association of the tRNA splicing endonuclease with the RNA kinase CLP1 ^5,22^. Mutations in CLP1 were shown to impair tRNA splicing *in vitro* and to cause neuropathologies involving the central and peripheral nervous system ^23–25^. Mutations in all four subunits of the TSEN complex have been associated with the development of pontocerebellar hypoplasia (PCH), a heterogeneous group of inherited neurodegenerative disorders with prenatal onset characterized by cerebellar hypoplasia and microcephaly ^26–31^.

High expression of TSEN54 in neurons of the pons, cerebellar dentate, and olivary nuclei suggested that a functional endonuclease complex is essential for the development of these regions ^26^. The most common mutation causing a type 2 PCH phenotype is a homozygous *TSEN54* c.919G>T mutation that leads to an A^307^S substitution in TSEN54 ^26,29^. Other substitutions, *e.g.* S^93^P in TSEN54, R^58^W in TSEN34, Y^309^C in TSEN2, and H^116^Y in TSEN15, have also been identified as causative for PCH ^26,27^. None of the described disease mutations are located in or in close proximity to the predicted catalytic sites of human TSEN, or in other highly conserved regions of the proteins, and how they contribute to disease development remains enigmatic. Here we present the biochemical and structural characterization of recombinant human TSEN. We analyze PCH-associated mutations at the structural and biochemical levels in reconstitution experiments and reveal biochemical features of the TSEN complex in PCH patient-derived cells.

## Results

### Assembly of recombinant human tRNA splicing endonuclease

To gain functional insights into human TSEN/CLP1, we designed an expression vector series based on the MultiBac system ^32^ that allows combinatorial protein complex production in insect and mammalian cells by utilizing a CMV/p10 dual promoter ^33^ (Fig. 1a,b and Extended Data Fig. 1a). Using this system, we were able to assemble and purify functional heterotetrameric TSEN and a heteropentameric complex including the RNA kinase CLP1 from infected insect cells (Fig. 1b,c and Extended Data Fig. 1b). The structural integrity of the purified complexes was verified by native mass spectrometry (MS), showing a stoichiometric TSEN2–15–34–54 heterotetramer (165.6 kDa) and a TSEN/CLP1 heteropentamer (213.0 kDa) (Fig. 1c, Extended Data Fig. 1b and Supplementary Tables 1,2). These data are in line with a recent study showing reconstitution of TSEN/CLP1 from a bacterial expression host ^34^. We also identified TSEN/CLP1 complexes harboring two CLP1 molecules (Extended Data Fig. 1b). Recombinant TSEN54 showed a high degree of phosphorylation as reported for the endogenous protein (Extended Data Fig. 1c) ^35^.

**Figure 1.**
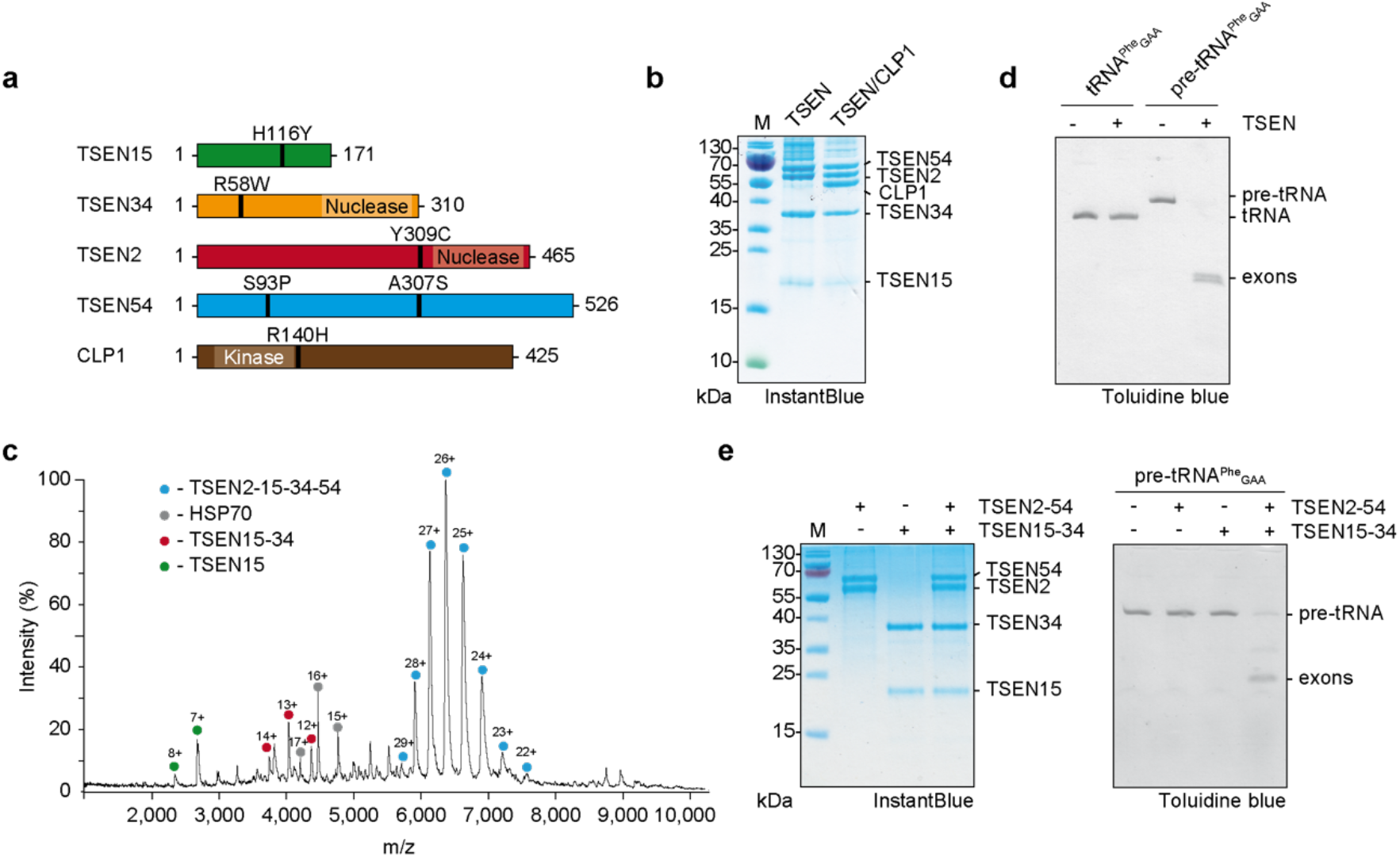
Assembly and catalysis of recombinant human TSEN. **a**, Bar diagrams of TSEN subunits and CLP1 depicting positions of PCH mutations, predicted nuclease domains of TSEN2 and TSEN34, and the RNA kinase domain for CLP1. Total amino acids of each protein are indicated. **b**, SDS-PAGE of purified recombinant TSEN and TSEN/CLP1 complexes visualized by InstantBlue staining. Protein identities and size markers are shown. **c**, Native mass spectrum of tetrameric TSEN complex from an aqueous ammonium acetate solution. Charge states of the predominant TSEN2–15– 34–54 heterotetramer (blue circles), Heat Shock Protein (HSP) 70 (grey circles), the heterodimer TSEN15–34 (red circles) and TSEN15 (green circles) are indicated. **d**, Pre-tRNA cleavage assay using tetrameric TSEN complex with *Saccharomyces cerevisia*e (*S.c.*) pre-tRNA^Phe^_GAA_ and mature tRNA^Phe^_GAA_. Input samples and cleavage products were separated via Urea-PAGE and visualized by Toluidine blue. RNA denominations are given on the right. **e**, Pre-tRNA cleavage assay with TSEN heterodimers and *S.c.* pre-tRNA^Phe^_GAA_. SDS-PAGE of the indicated heterodimers and the reconstituted TSEN tetramer is shown on the left (InstantBlue stain), Urea-PAGE of the cleavage products on the right (Toluidine blue stain). Gels are representative of three independent experiments. Unprocessed gels for **b**, **c** and **e** are shown in Source Data 1.

Endonuclease activity of tetrameric TSEN was observed in a pre-tRNA cleavage assay using *Saccharomyces cerevisiae* (*S.c.*) pre-tRNA^Phe^_GAA_ as a substrate, whereas mature *S.c.* tRNA^Phe^_GAA_ remained unaffected (Fig. 1d). The absence of endonucleolytic activity on mature tRNA confirms the specificity of the complex for its native pre-tRNA substrate. Yeast-two-hybrid experiments with *S.c.* orthologues suggested that strong interactions exist between TSEN15 and TSEN34, as well as between TSEN2 and TSEN54, and that the human endonuclease assembles from preformed dimeric subcomplexes ^8^. Using combinatorial co-expression analyses, we identified the formation of stable TSEN15–34 and TSEN2–54 heterodimers (Fig. 1e). Individual heterodimers did not show endonuclease activity, whereas specific endonucleolytic cleavage was observed after stoichiometric mixing of TSEN15–34 and TSEN2–54 in the absence of ATP (Fig. 1e). Size exclusion chromatography confirmed that a stable tetrameric assembly formed upon mixing the individual heterodimers (Extended Data Fig. 1d,e). These observations indicate that active human TSEN assembles from non-functional, heterodimeric submodules independently of CLP1 and ATP.

### Human TSEN binds precursor and mature tRNAs with similar affinities

It has been postulated that eukaryotic splicing endonucleases recognize pre-tRNAs via their mature domain ^14,15^. To define tRNA binding parameters of human TSEN, we performed interaction studies using catalytically inactive tetramers (TSEN^*inactive*^), in which the active site histidines of TSEN2 (His^377^) and TSEN34 (His^255^) were substituted with alanines (Fig. 2 and Extended Data Fig. 2). Alanine substitutions of His^377^ of TSEN2 and His^255^ of TSEN34 abolished cleavage at the 5’ and 3’ splice sites, respectively, and purified TSEN^*inactive*^ did not cleave pre-tRNA substrates at all (Fig. 2a and Extended Data Fig. 2b).

**Fig. 2.**
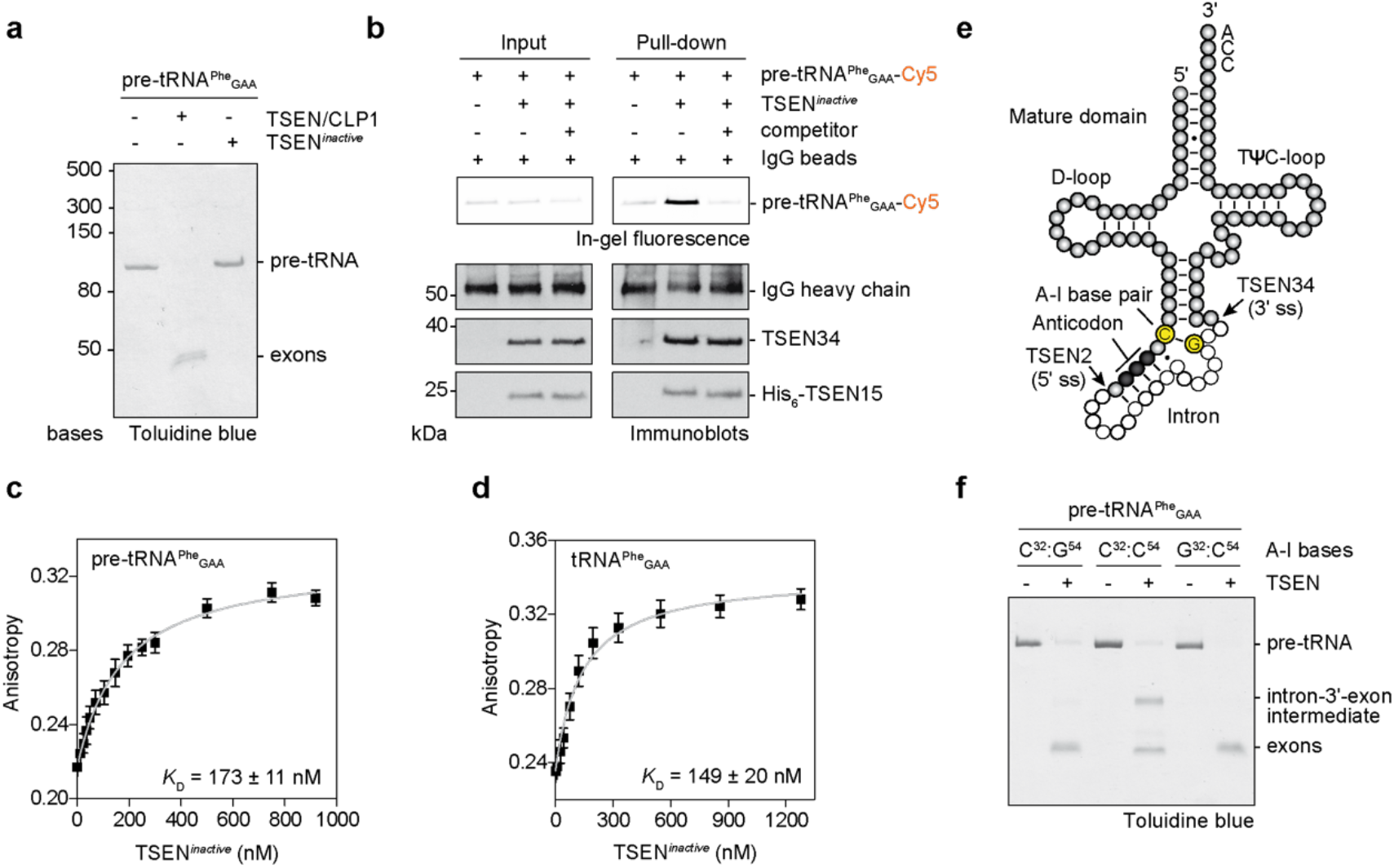
Active involvement of the A–I base pair in coordinating pre-tRNA cleavage. **a**, Pre-tRNA cleavage assay comparing recombinant, inactive TSEN tetramer (TSEN2^H377A^ and TSEN34^H255A^ double mutant) to the TSEN/CLP1 complex. Cleavage products are visualized by denaturing Urea-PAGE with subsequent Toluidine blue staining. RNA size markers are indicated on the left of the gel. **b**, Pull-down assay with fluorescently labeled *S.c.* pre-tRNA^Phe^_GAA_ and inactive, tetrameric TSEN captured on protein G agarose functionalized with an α-His-antibody. Protein size markers are indicated on the left of each immunoblot, protein and RNA identities on the right. Input and co-precipitated, labeled pre-tRNAs were visualized by in-gel fluorescence, TSEN subunits and the immunoglobulin G (IgG) heavy chain by immunoblotting. The IgG heavy chain served as loading control. **c,** Thermodynamic binding parameters of fluorescently labeled *S.c.* pre-tRNA^Phe^_GAA_ and inactive, tetrameric TSEN revealed by fluorescence anisotropy. **d**, Thermodynamic binding parameters of fluorescently labeled tRNA^Phe^_GAA_ and inactive, tetrameric TSEN revealed by fluorescence anisotropy. **e**, Schematic depiction of a pre-tRNA molecule showing ribonucleotides belonging to the mature domain (grey spheres), the intronic region (white spheres), the anticodon (black spheres), and the A-I base pair (yellow spheres). Proposed 5’ and 3’ splice sites (ss) are indicated. **f**, Impact of A-I base pair mutations in *S.c.* pre-tRNA^Phe^_GAA_ on endonucleolytic activity of tetrameric TSEN revealed by a pre-tRNA cleavage assay. C^32^:G^54^ – canonical A-I base pair, C^32^:C^54^ – disrupted A-I base pair, G^32^:C^54^ – inverted A-I base pair. All experiments are representatives of three independent assays. Unprocessed gels for **a**, **b** and **f** are shown in Source Data 3.

To perform fluorescence anisotropy and pull-down experiments, we site-specifically labeled precursor and mature yeast tRNA^Phe^_GAA_ at the terminal 3’ ribose. Despite the inability to cut its native substrate, TSEN^*inactive*^ interacted stably and specifically with the fluorescently labeled pre-tRNA in a pull-down assay (Fig. 2b). Binding studies using fluorescence anisotropy revealed dissociation constants (*K*_*D*_) of 173±11 nM and 149±20 nM for fluorescently labeled pre-tRNA^Phe^_GAA_ and mature tRNA^Phe^_GAA_, respectively (Fig. 2c,d). We determined an inhibition constant (*K*_i_) of 197 nM (95% confidence interval of 168 – 231 nM) in a competition assay confirming the specific interaction, whereas a fluorescent electrophoretic mobility shift assay corroborated a dissociation constant between TSEN and pre-tRNA in the high nanomolar range (Extended Data Fig 2c,d). Our determined *K*_D_ values are in good agreement with previously deduced Michaelis constants (*K*_*M*_) of ~30 nM and 250 nM for intron excision by the yeast and an archaeal tRNA endonuclease, respectively ^36^. Taken together, the results show that substrate recognition by human TSEN is primarily mediated by the mature domain of pre-tRNAs but not their introns and suggests that discrimination between pre- and mature tRNAs might be dictated by kinetic effects.

### The A–I base pair coordinates cleavage at the 3’ splice site in human TSEN

Cleavage of archaeal introns strictly relies on the tRNA BHB motif, whereas the only preserved feature of human tRNA introns is a pyrimidine in the 5’ exon at position −6 from the 5’ splice site which forms a conserved A–I base pair with a purine base at position −3 from the 3’ splice site (Fig. 2e). Studies on the *Xenopus* tRNA endonuclease showed that the A–I base pair is critically involved in the cleavage reaction at the 3’ splice site ^20,21,37^. To find out whether the same regulatory principles exist for intron excision in humans, we tested the impact of A–I base pair mutants on endonucleolytic cleavage by tetrameric TSEN (Fig. 2e,f, Extended Data Fig. 2e, and Supplementary Fig. 1). Changing the guanine base G^54^ to cytosine in *S.c.* pre-tRNA^Phe^_GAA_ produced a pre-tRNA substrate with a disrupted A–I base pair (Fig. 2e,f). Cleavage of this pre-tRNA by wild-type (wt) human TSEN resulted in a 5’ exon and an intron-3’-exon intermediate (Fig. 2f). Cleavage at both splice sites was observed when base pairing at the A–I position was restored by mutating cytosine C^32^ to guanine in the C^54^ background (Fig. 2e,f). The same effect was observed when human pre-tRNA^Tyr^_GTA_8-1 harboring equivalent mutations was used as substrate (Extended Data Fig. 2e). These findings imply that the presence of an A–I base pair, but not the strict identity of the bases, is essential for cleavage at the 3’ splice site by human TSEN ^20,21^.

### The molecular architecture of TSEN is evolutionarily conserved

Our interaction studies using recombinant proteins showed that active human TSEN assembles from inactive TSEN15–34 and TSEN2–54 heterodimers (Fig. 1e). To gain detailed insights into the molecular architecture of the human TSEN complex, we characterized the TSEN15–34 heterodimer by X-ray crystallography (Fig. 3 and Extended Data Fig. 3). Despite extensive crystallization trials, full-length TSEN15–34 did not crystallize. To define a crystallizable core complex, we subjected the TSEN15–34 complex to limited proteolysis with subsequent size exclusion chromatography and MS analysis (Extended Data Fig. 3a,b and Supplementary Tables 2-4). We observed two comigrating polypeptide species corresponding to residues 23 to 170 of TSEN15 and residues 208 to 310 of TSEN34 covering the predicted conserved nuclease domains (Extended Data Fig. 3b,c and Supplementary Fig. 2) ^8^.

**Fig. 3.**
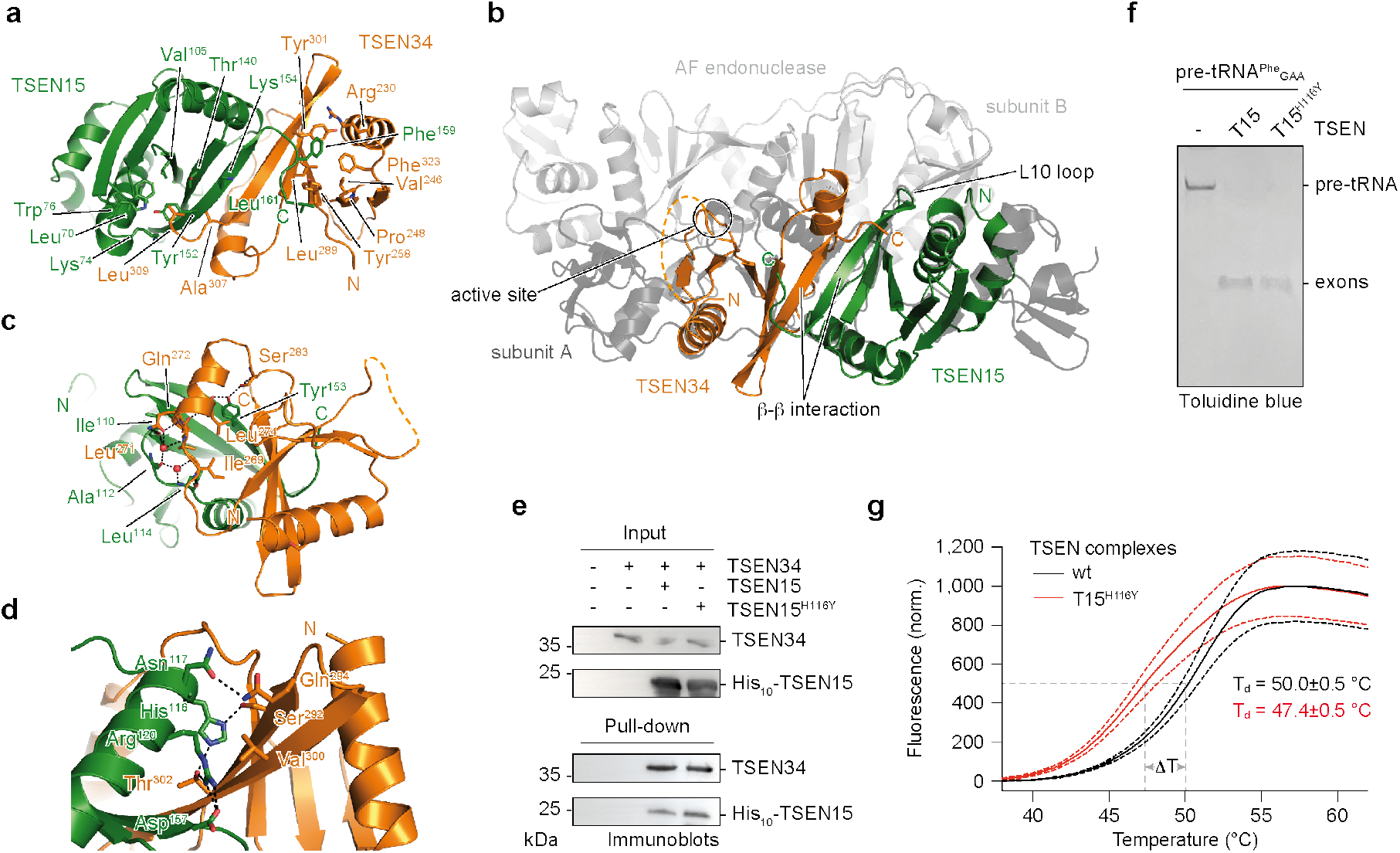
Structural and functional details of the TSEN15–34 dimer interface. **a**, X-ray crystal structure of a TSEN15–34 complex derived from limited proteolysis experiments. TSEN15 (green) and TSEN34 (orange) are shown in cartoon representation. Key amino acids are depicted in stick representation together with amino-(N) and carboxy (C)-termini. **b**, Superposition of the TSEN15–34 heterodimer and the pre-tRNA endonuclease from *Archaeoglobus fulgidus* (AF) (PDB ID 2GJW) ^13^. The position of the catalytic triad of TSEN34 (active site), the L10 loop of TSEN15 and the β-strands involved in the β-β interaction between TSEN15 and TSEN34 are shown. **c**, Cartoon representation of the dimer interface with amino acid residues in stick representation (color coding as in **a**). Water molecules and hydrogen bonds are shown as red spheres and black dashed lines, respectively. **d**, Cartoon representation of the TSEN15–34 interface highlighting histidine 116 (His^116^) of TSEN15, mutated in patients with a PCH type 2 phenotype. **e**, Pull-down experiment with TSEN34, wt TSEN15 and TSEN15 carrying the H116Y mutation. Input and co-precipitated proteins were separated by SDS-PAGE and visualized by immunoblotting. Size markers and protein identities are shown. **f**, Pre-tRNA cleavage assay with wt, tetrameric TSEN complex and a tetrameric TSEN complex carrying the TSEN15^H116Y^ (T15^H116Y^) mutation. Cleavage products were separated by Urea-PAGE and visualized with toluidine blue. **g**, Thermal stability of wt, tetrameric TSEN (black line) and TSEN15^H116Y^ mutant complex (red line) assessed by differential scanning fluorimetry (DSF). Note that recombinant complexes were purified from HEK293 cells. Normalized (norm.) fluorescence is plotted against temperature in degree Celsius (°C). Denaturation temperatures (T_*d*_) of the complexes were derived from sigmoidal Boltzmann fits (grey dashed lines) with error of fits. Standard deviations (SD) of technical triplicates are represented by red (T15^H116Y^) and black (wt) dashed lines. Unprocessed gels for **e** and **f** are shown in Source Data 5.

We re-cloned, co-expressed and purified the proteolytically characterized fragments, which readily formed rod-shaped crystals in space group P2_1_ and diffracted X-rays to a resolution of 2.1 Å (Extended Data Fig. 3d and Supplementary Table 5). The asymmetric unit is composed of two domain-swapped TSEN34 molecules, each binding one TSEN15 protomer at their C-terminal domains (Extended Data Fig. 3e,f). The domain swap is brought about by a short, structured N-terminal α-helix/β-hairpin element of TSEN34 that is liberated to hook onto the neighboring protomer, presumably due to the truncated N-terminus of the molecule. The domain-swap organization is only found under crystallization conditions, as shown by size exclusion chromatography multi-angle light scattering (SEC-MALS) (Extended Data Fig. 3g). The two TSEN15 and the two TSEN34 molecules in the asymmetric unit are very similar with average overall RMSDs of 0.37 Å and 0.50 Å, respectively. In one TSEN15 protomer, an elongated N-terminal region (residues 162-170) is visible, which is stabilized by crystal contacts (Extended Data Fig. 3f).

TSEN15 and TSEN34 display the typical endonuclease fold with the latter harboring the Tyr^247^/His^255^/Lys^286^ catalytic triad as also found in archaeal and eukaryotic endonucleases (Fig. 3a,b and Extended Data Fig. 3h) ^13^. The dimeric TSEN15–34 complex is characterized by an elongated central twisted β-sheet connected by the C-terminal β-strands of TSEN15 and TSEN34 with a buried surface area of ~ 1980 Å^2^ between the protomers (Fig. 3a,b). Each face of the individual twisted β-sheets of TSEN15 and TSEN34 is mainly stabilized by hydrophobic interactions to an alpha helix (Fig. 3a,c,d). In the interface between TSEN15 and TSEN34 two structural water molecules are found, which are coordinated by hydrogen bonds to backbone oxygens or amide groups of Ile^110^, Ala^112^, and Leu^114^ of TSEN15, Ile^269^, Leu^271^, and Gln^272^ of TSEN34, and the side chain oxygen of Gln^272^ (Fig. 3c). Furthermore, a YY motif in TSEN15 (Tyr^152^/Tyr^153^), which is conserved in eukaryotic endonucleases and archaeal α_4_- and (αβ)_2_-type endonucleases (Supplementary Fig. 3) both stabilizes TSEN15 by hydrophobic interactions and the dimer interface by hydrogen bonds to the main chain carbonyl oxygen of Leu^274^ and the side chain oxygen of Ser^283^ of TSEN34 (Fig. 3a,c). In contrast to a previous solution NMR structure of homodimeric TSEN15 ^9^, our biochemical and structural analyses show that the assembly and architecture of TSEN are conserved and support the hypothesis that tRNA splicing endonucleases arose from a common ancestor through gene duplication and differentiation events ^38^.

### PCH-causing mutations destabilize recombinant TSEN

A previous genetic study identified a His-to-Tyr substitution at position His^116^ of TSEN15 in patients with PCH type 2 (Fig. 3d) ^27^. The imidazole group of His^116^ is central to a hydrogen bond network involving Asn^117^, Arg^120^, and Asp^157^ of TSEN15 and Ser^292^ and Thr^302^ of TSEN34 (Fig. 3d). We tested the impact of this substitution in a pull-down experiment using full-length TSEN15 and TSEN34 and also in a pre-tRNA cleavage assay in the context of the tetrameric assembly (Fig. 3e,f). We hypothesized that the substitution impairs complex formation and activity due to steric clashes in the dimer interface and loss of the hydrogen bond network. However, TSEN15 carrying the His-to-Tyr mutation engaged in complex formation with TSEN34 similar to the wt protein, and no impairment of catalytic activity was observed (Fig. 3e,f). We assumed that the large hydrophobic interface compensates for the loss of the hydrogen bond network. To assess the effects of the TSEN15^H116Y^ mutation on the thermal stability of TSEN, we compared the mutant complex to wt by differential scanning fluorimetry (DSF, Fig. 3g) ^39^. This assay reported apparent denaturing temperatures of 50.0±0.5 °C and 47.4±0.5 °C for wt and mutant TSEN, respectively (Fig. 3g and Extended Data Fig. 3i). These data suggest that destabilization of TSEN might be a general effect of PCH-causing mutations.

The molecular basis of PCH mutations on disease development is largely unknown ^30^. It is hypothesized that mutations in TSEN contribute to the disease by interfering with complex assembly, stability, or enzymatic activity. Given that the His-to-Tyr mutation in TSEN15 thermally destabilized the endonuclease complex, we produced heterotetrameric TSEN complexes carrying the PCH-causing mutations TSEN2^Y309C^, TSEN34^R58W^, TSEN54^S93P^ and TSEN54^A307S^ in HEK293 cells and performed pull-down experiments to assess complex assembly and integrity (Fig. 4a). Despite subtle differences in expression levels of the individual subunits, pull-down via TSEN15 co-precipitated TSEN2, TSEN34, and TSEN54, irrespective of the introduced PCH-causing mutation (Fig. 4a). Control pull-downs from HEK293 cells overexpressing only His-tagged TSEN15 showed that endogenous subunits of TSEN do not associate with overexpressed TSEN15, probably due to their very low copy numbers (Extended Data Fig. 4a). We produced and purified recombinant heterotetrameric TSEN complexes carrying the pathogenic missense mutations from baculovirus-infected insect cells (Fig. 4b) and also did not see obvious deleterious effects on subunit composition or pre-tRNA cleavage kinetics (Fig. 4b,c and Extended Data Fig. 4b).

**Fig. 4.**
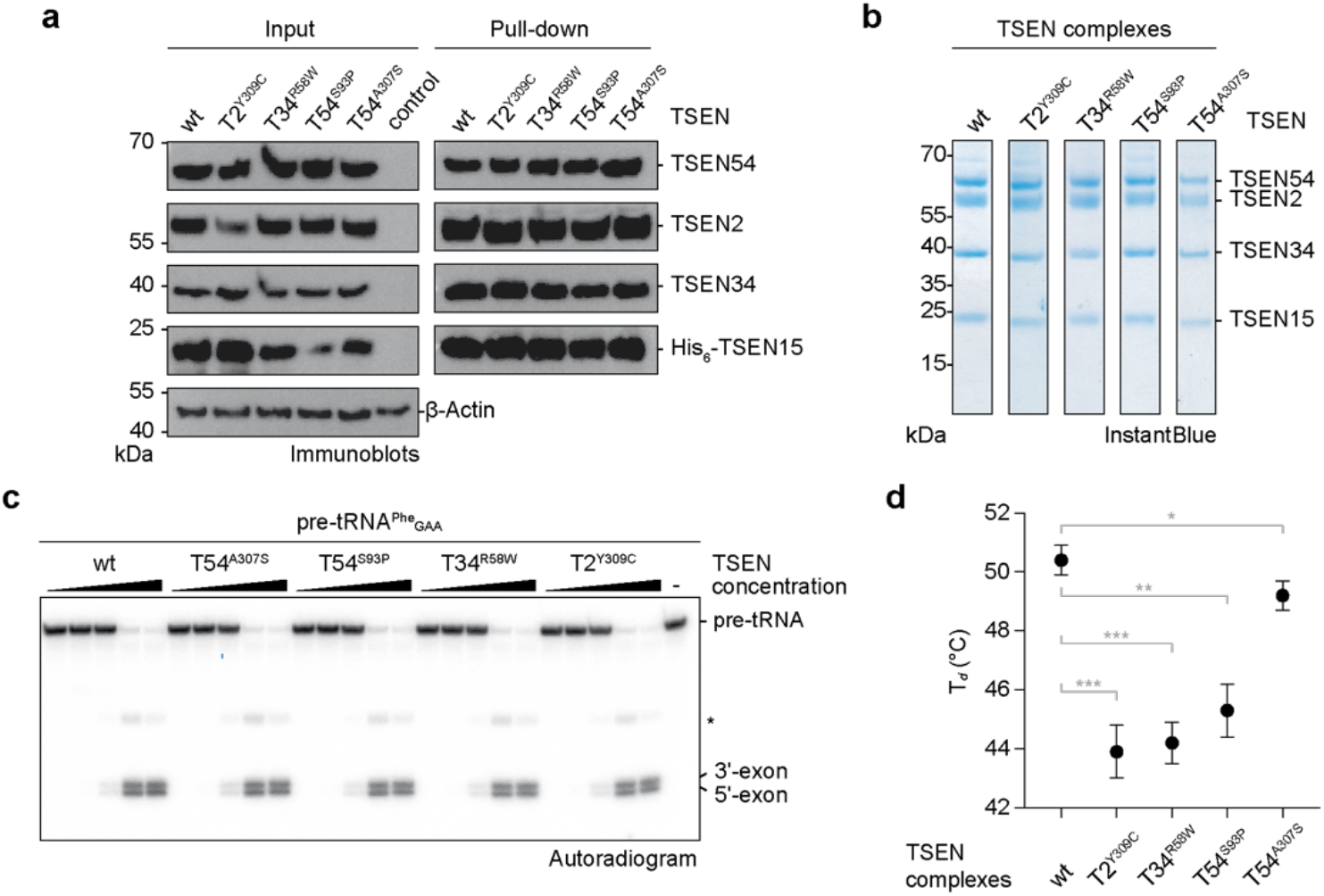
PCH mutations affect thermal stability but not activity of recombinant TSEN. **a**, Immunoblot analysis of a pull-down assay with wt and mutant TSEN complexes co-expressed in HEK293 cells. Size markers and protein identities are indicated. **b**, SDS-PAGE of SEC peak fractions of purified recombinant wt and mutant heterotetrameric TSEN complexes. **c**, Pre-tRNA endonuclease assay of radioactively labelled *S.c.* pre-tRNA^Phe^_GAA_ with increasing amounts of recombinant wt or mutant TSEN complexes revealed by phosphorimaging. **d**, Thermal stability of recombinant wt and mutant TSEN complexes assessed by DSF. Shown is the denaturation temperature (T_*d*_) of each complex with fit errors derived from Boltzmann sigmoids. Fit errors were derived from means of technical triplicates and are representative of biological duplicates. Unprocessed gels for **a**, **b**, and **c** are shown in Source Data 7 and 8.

Given the low abundance of TSEN molecules in cells ^8^ and that PCH mutations phenotypically affect only cerebellar neurons, we reasoned that expression levels are too high in our reconstitution systems to reveal subtle alterations in TSEN assembly and function. To assess the effects of PCH-causing mutations on complex stability, we used the DSF assay (Fig. 3g and Extended Data Fig. 3i). Most PCH-causing mutations led to substantial shifts towards lower denaturation temperatures (*e.g.* T_*d*_ of 43.9±0.9 °C for TSEN2^Y309C^ compared to T_*d*_ of 50.4±0.5 °C for wt TSEN) when exposed to thermal gradients indicating that mutant TSEN complexes have compromised structural integrity (Fig. 4d and Extended Data Fig. 4c). The relative changes in thermostability (T_Δ_) of the mutant complexes compared to wt TSEN ranged from 6.5 °C for the TSEN2^Y309C^ mutation to 1.2 °C for the TSEN54^A307S^ mutation potentially scaling with the severity of the disease phenotype (Fig. 4d and Extended Data Fig. 4c) ^26^. DSF data also revealed two-state unfolding behaviors for all TSEN complexes when analyzed by the ProteoPlex algorithm ^40^ suggesting cooperativity of unfolding transitions for the individual subunits (Supplementary Table 6), thus explaining why mutations in different subunits lead to an overall destabilization of TSEN. Our data suggest that PCH phenotypes in patients potentially develop due to destabilized TSEN complexes.

### Pre-tRNA processing is impaired in PCH patient cells

To determine if pre-tRNA processing activity is compromised in PCH patients we derived primary skin fibroblasts from PCH patients, their healthy parents, and unrelated controls (Supplementary Table 7). We chose the common *TSEN54* c.919G>T (TSEN54^A307S^) mutation, which is reported in ~ 90% of recognized TSEN-linked PCH cases ^30^, and for which a large cohort of patient samples are available. The cell lines we created did not show any morphological differences compared to control cells. When we assayed lysates derived from homozygous *TSEN54* c.919G>T cell lines, we observed a reduction in pre-tRNA splicing efficiency compared to control cell lysates (Fig. 5a). Subtle differences in ligation efficiency, as observed for cell line Ba2, may result from the fibroblasts having different genetic backgrounds.

**Fig. 5.**
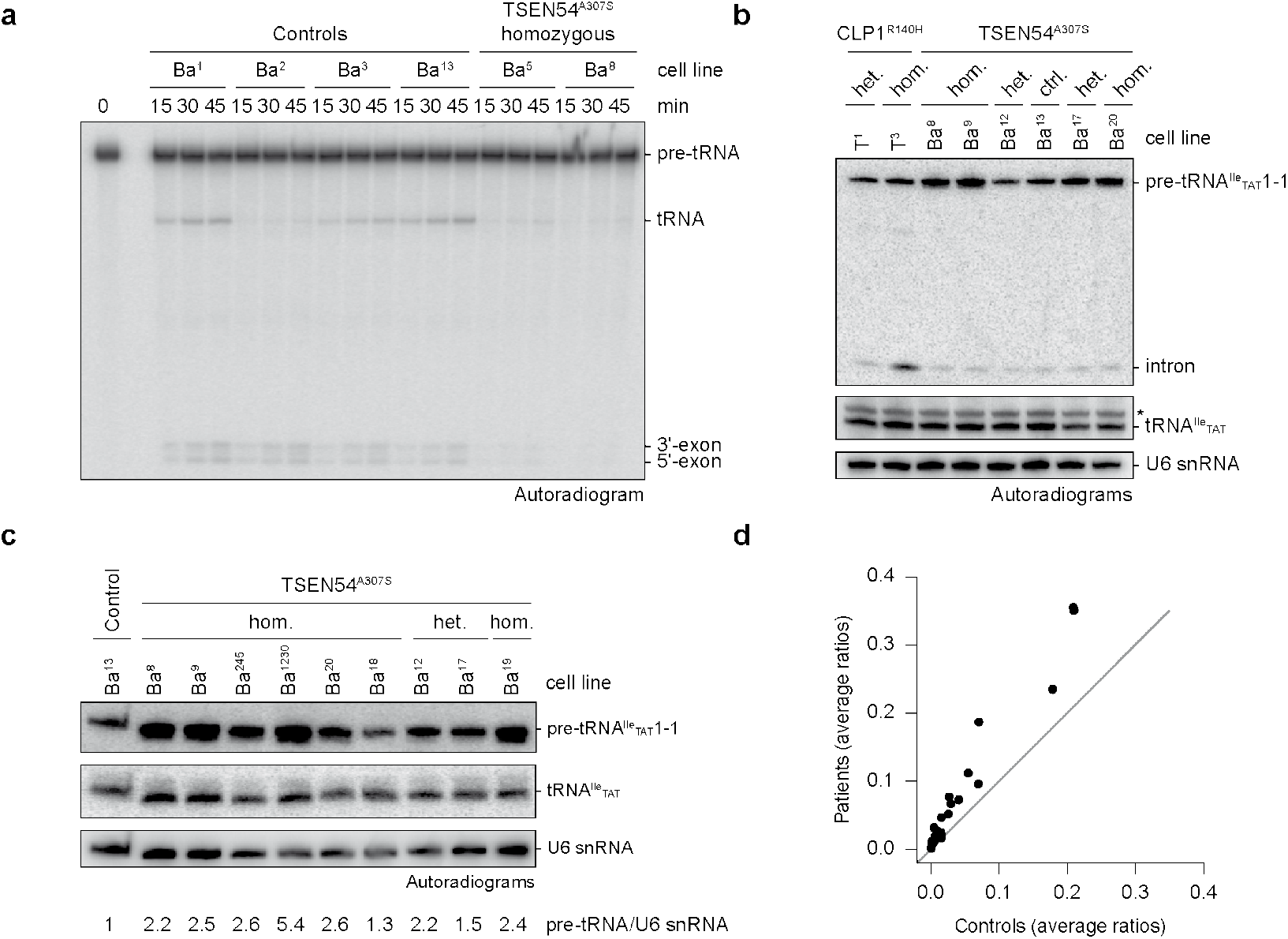
*TSEN54* c.919G>T fibroblasts exhibit reduced splicing activity *in vitro* and accumulation of intron-containing pre-tRNAs. **a**, Pre-tRNA splicing assay (time course) with radioactively labelled *S.c.* pre-tRNA^Phe^_GAA_ and cell extracts derived from control and PCH patient fibroblasts. Splice products were separated by Urea-PAGE and monitored by phosphorimaging. **b**, Comparison of pre-tRNA^Ile^_TAT_1-1 intron abundance between control cells and fibroblasts carrying the heterozygous or homozygous *TSEN54 c.919*G>T (TSEN54^A307S^) or the *CLP1 c.419*G>A (CLP1^R140H^) mutation by northern blotting. **c**, Northern blot analysis comparing pre-tRNA^Ile^_TAT_1-1 and mature tRNA^Ile^_TAT_1-1 levels to levels of U6 snRNA in control fibroblasts and fibroblast carrying the heterozygous (het.) or homozygous (hom.) *TSEN54 c.919*G>T mutation with quantification. **d**, Average ratios of pre-tRNAs to mature tRNAs derived from Hydro-tRNAseq for all intron-containing tRNAs comparing PCH patients to control samples. The black line indicates a slope of 1. Panels in **a**, **b**, and **c**, are representative of at least two independent experiments. Unprocessed gels for **a**, **b**, and **c**, are shown in Source Data 10.

This result is reminiscent of observations in patient-derived cell lines carrying a homozygous *CLP1* c.419G>A (CLP1^R140H^) mutation, which leads to severe motor-sensory defects, cortical dysgenesis, and microcephaly ^23,25^. In contrast to homozygous CLP1^R140H^ cells, in which introns accumulate, intron accumulation did not occur in either heterozygous or homozygous TSEN54^A307S^ backgrounds as judged by northern blot analysis using a probe specific for the intron of pre-tRNA^Ile^_TAT_1-1 (Fig. 5b). These results suggest an impairment of intron excision rather than a defect in downstream processes of the tRNA splicing reaction, which might lead to the accumulation of pre-tRNAs in patient cells.

To test this hypothesis, we compared levels of intron-containing pre-tRNAs to their corresponding mature tRNAs in cell lines carrying the homozygous *TSEN54* c.919G>T mutation to heterozygous cell lines and controls by hydro-tRNAseq ^4^ and northern blotting (Fig. 5c,d, Extended Data Fig. 5, and Supplementary Table 8). We observed an accumulation (~2-6 fold) of intron-containing pre-tRNAs in homozygous *TSEN54* c.919G>T cell lines compared to control cell lines, albeit global levels of mature tRNAs remained largely unaffected (Fig. 5c,d and Extended Data Fig. 5). The distributions of the ratios of precursor over mature tRNA reads showed that there was no bias for an enrichment of a specific precursor tRNA among samples (Extended Data Fig. 5a and Supplementary Table 8). These findings are consistent with the observation that homozygote patient samples exhibited a relative increase in precursor tRNA reads, compared to wild-type controls (Fig. 5d and Extended Data Fig. 5b). Therefore, we conclude that in our experimental setup, TSEN54 A^307^S results in an increase of the steady-state levels of intron-containing tRNAs. Consistently, northern blot analyses showed similar differences in levels of pre-tRNA^Ile^_TAT_1-1 over mature tRNA^Ile^_TAT_1-1 (Fig. 5c). Taken together, our data show defects in pre-tRNA processing uncoupled from CLP1 function leading to the accumulation of pre-tRNAs in PCH patient-derived cell lines.

### Pre-tRNA processing defects are linked to altered TSEN composition

To investigate whether the reduction of pre-tRNA processing in cell extracts of homozygous *TSEN54* c.919G>T patients was due to altered TSEN assembly or stability we used rabbit polyclonal antibodies against peptides of TSEN2, TSEN34, and TSEN54 ^35^, to asses changes in TSEN subunit abundance and to perform co-immunoprecipitation experiments of endogenous TSEN. Immunoblot analyses showed that the homozygous *TSEN54* c.919G>T mutation does not impact steady-state levels of TSEN54, suggesting that no changes in either mRNA stability, transcription rate or protein turnover occur (Fig. 6a).

**Fig. 6.**
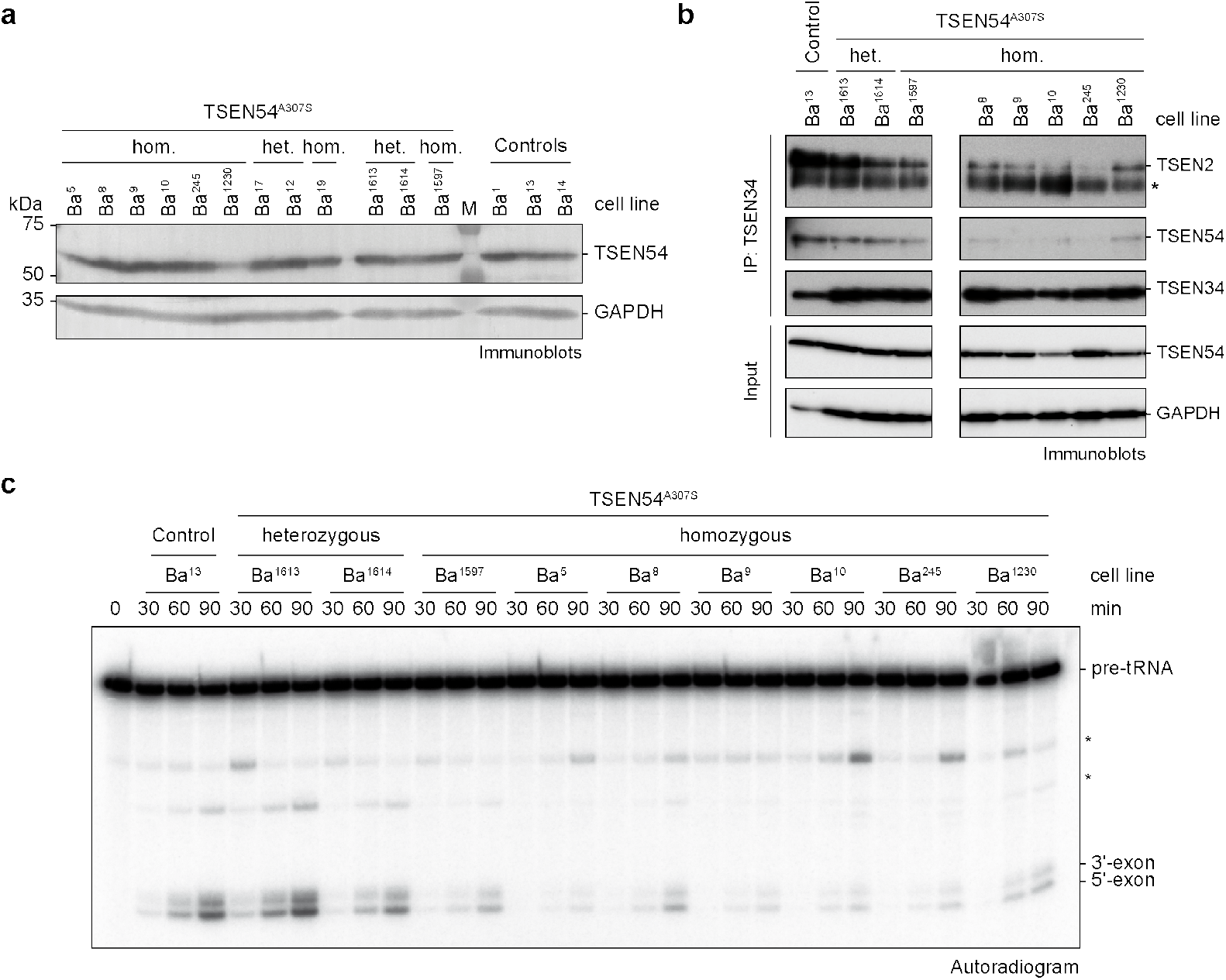
Reduced pre-tRNA cleavage activity in PCH patient-derived cell extracts is associated with altered composition of TSEN. **a**, Comparison of TSEN54 protein levels between control and heterozygous or homozygous PCH patient fibroblasts analyzed by immunoblotting. GAPDH served as a loading control. **b**, Co-immunoprecipitation (IP) assay using an α-TSEN34 antibody with cell lysates derived from control and heterozygous or homozygous *TSEN54 c.919*G>T fibroblasts analyzed by immunoblotting. The asterisk indicates the heavy chain of the α-TSEN34 antibody. **c**, On-bead pre-tRNA cleavage assay (time course) with radioactively labelled *S.c.* pre-tRNA^Phe^_GAA_ and immunoprecipitated TSEN complexes (α-TSEN34 antibody-coupled resin) derived from control fibroblasts and from fibroblasts carrying heterozygous or homozygous *TSEN54 c.919G*>T mutation shown in (**b**). Unspecific bands are indicated by asterisks. Data are representative of at least two independent experiments. Unprocessed gels for **a**, **b**, and **c** are shown in Source Data 11-13.

To evaluate TSEN complex composition and pre-tRNA cleavage activity we performed immunoprecipitation experiments from patient-derived and control fibroblasts using α-TSEN2 or α-TSEN34 antibodies (Fig. 6b,c and Extended Data Fig. 6). Immunoblot analyses showed a substantial reduction of co-immunoprecipitated TSEN2 and TSEN54 from patient cell lines using an α-TSEN34 antibody, while at the same time, pre-tRNA cleavage activity was strongly diminished in α-TSEN2 and α-TSEN34 immunoprecipitates (Fig. 6b,c and Extended Data Fig. 6). These results indicate that TSEN assembly defects lead to reduced pre-tRNA cleavage in PCH patient cells. Since the association of TSEN2 and TSEN54 is likewise affected but steady-state levels of the individual proteins are not, we conclude that impaired TSEN activity is caused by an altered propensity for the formation of the active tetrameric assembly in patient cells.

## Discussion

Here we report the recombinant expression, purification, and assembly of functional human TSEN/CLP1 complex. We show that heterotetrameric TSEN is assembled from heterodimeric TSEN15–34 and TSEN2–54 subcomplexes, which combine to form the composite active sites for catalysis (Fig. 1e). The nuclease fold seen in our TSEN15–34 X-ray crystal structure is conserved with the archaeal tRNA endonucleases (Fig. 3b) ^13,16^ suggesting that the TSEN2–54 heterodimer — and entire TSEN complex — likely forms through interactions similar to those seen in the TSEN15–34 heterodimer, as well as related interactions previously observed in archaeal tRNA endonucleases. Our interaction studies with catalytically inactive TSEN mutants show that substrate recognition occurs through interactions with the mature tRNA fold, including the aminoacyl acceptor stem, the D-loop, and the Ψ-loop, and support the ‘ruler model’ of substrate recognition (Fig. 2c,d) ^8^. The similar affinities TSEN shows for pre-tRNAs and tRNAs suggest thermodynamic effects are unlikely to play a role in substrate selection (Fig. 2c,d). Instead, we speculate that different binding kinetics should contribute to the selection of pre-tRNAs over mature tRNAs, thereby guaranteeing efficient scanning and processing of the large pre-tRNA pool.

The tRNA splicing machinery is involved in processing of other RNA species ^41–44^. Eukaryotic tRNA endonucleases are involved in processing of mRNAs and rRNAs ^5,44,45^. TSEN is a key factor in the generation of tRNA intronic circular (tric) RNAs, a poorly uncharacterized class of short non-coding RNAs in *Drosophila* and humans ^41^. Archaeal tRNA endonucleases are capable of binding and cutting any RNA fragment that adopts a BHB motif ^19^. tRNA splicing in *Xenopus* necessitates a purine/pyrimidine base pair at the A–I base pair positions for 3’ splice site recognition and cleavage. Our data show that requirements for cleavage at the 3’ splice site by human TSEN are more relaxed and only need the A–I base pair, whereas the purine/pyrimidine identities of the bases are negligible (Fig. 2f and Extended Data Fig. 2e). The relaxed specificity may facilitate recognition and cleavage of non-canonical substrates. However, structures of human tRNA endonucleases with bound pre-tRNA substrate confirming this hypothesis are still missing. Nonetheless, our data suggest that substrate recognition and cleavage by human TSEN are two distinct processes with different structural requirements regarding the RNA.

While we show that assembly and enzymatic function of recombinant human TSEN complexes are immune to PCH-associated mutations, these mutations cause thermal destabilization with apparent deleterious effects on complex assembly and activity in patient cells (Fig. 3e,f,g, Fig. 4, Extended Data Fig. 4, Fig. 5, Extended Data Fig. 5, Fig. 6, Extended Data Fig. 6). Structural studies on archaeal tRNA endonucleases show that there are two major interaction interfaces: The β-β-interaction, mainly driven by hydrophobic interactions, and the L10 loop, involving hydrogen bonds and salt bridges. The hydrophobic interface has a higher degree of plasticity and thereby could accommodate mutations to a certain extent, whereas interactions within the hydrophilic interface are less tolerant of changes. Since TSEN is low-abundant (~ 100 molecules per cell) ^7^, destabilization by PCH-associated mutations may, therefore, have a strong effect on the assembly of the heterotetramer, whereas the individual heterodimers are sufficiently stable to escape protein degradation. In line with this hypothesis, we find decreased levels of TSEN2 and TSEN54 in α-TSEN34 immunoprecipitates from PCH patient cells (Fig. 6b and Extended Data Fig. 6a).

The question remains why TSEN mutations lead to a disease phenotype only in a subset of neurons, resulting in selective degeneration of cerebellar and, to a variable extent, anterior cortical structures ^29,46^. Neuropathologies caused by ablation of TSEN54 are not restricted to humans since knockdown of TSEN54 leads to brain hypoplasia in zebrafish ^46^ and a causative mutation in TSEN54 was identified in standard Schnauzers dogs with leukodystrophy ^47^.

Defective tRNA processing has been linked to various neuronal diseases ^48^. Although adequate supply of faithfully spliced tRNAs is expected to be essential for protein biosynthesis in all cell types ^49^, neurons may be particularly susceptible to subtle translation defects and, consequently, defects in proteostasis ^50^. Such delicate fine-tuning of translation is highly sensitive to changes in tRNA levels, which may influence the local speed of mRNA translation in a tissue-specific manner depending on the availability of cognate tRNAs ^51^. Balanced kinetics of tRNA accumulation could be crucial in tissues or cell sub-populations with a high metabolism, so that an otherwise modest defect in production rate might be deleterious where there is a high demand. Neurons require rapid and local protein synthesis for synaptic plasticity, which needs coordinated transport of the translational machinery, mRNAs, and tRNAs themselves. In line with these notions, missense mutations in subunits of the catalytic core of Pol III have been shown to affect assembly of the polymerase and have been linked to leukodystrophies ^52^. A large number of human neurodegenerative disorders have been linked to mutations in components of the general translational machinery and to numerous genes involved in tRNA expression and processing ^1,53^. With CLP1 ^23–25^ and arginyl tRNA synthetase ^54^ at least two other tRNA processing factors are linked to PCH ^29^ with only mild biochemical effects. Our data suggest that tRNA processing defects caused by TSEN or CLP1 mutations are distinct from one another, acting at different steps of splicing ^34^.

Impaired TSEN function may selectively impact the processing of cerebellum-specific pre-tRNAs. In mammals, expression of tRNA isoacceptor families (tRNAs with the same anticodon) varies between tissues and during development, and can be altered under disease conditions ^55–57^. Changes in tRNA repertoires have been hypothesized to correlate with the codon usage of genes associated with cellular differentiation states to fine-tune their translation ^55–57^. A mutation in a tRNA gene specifically expressed in the central nervous system has been shown to exhibit a synthetic effect with the loss of a ribosome recycling factor, selectively inducing cerebellar neurodegeneration in mice ^58^. In a similar scenario, neuron-specific isodecoders could be critically reduced in PCH patients, as a result of TSEN failure to cleave specific precursors.

tRNAs also function as signaling molecules in the regulation of numerous metabolic and cellular processes, or as stress sensors and in tRNA-dependent biosynthetic pathways ^59^. Transfer RNA-derived fragments (tRFs) have been identified as small non-coding RNAs contributing to translational control, gene regulation and silencing, as well as progressive motor neuron loss ^60^. Therefore, impaired TSEN activity could potentially result in unbalanced tRF levels with deleterious cellular effects.

While our data link a pre-tRNA splicing defect to PCH, additional factors and cellular mechanisms could be involved in the disease. Altered complex stability might affect interactions between TSEN and other cellular components. In yeast, TSEN activity has been linked to pre-rRNA and mRNA processing ^43,45^, thus certain neuron-specific mRNA transcripts might require some thus far uncharacterized activity of TSEN, which is impaired by the disease mutations. Clearly, future studies will be needed to address these questions *in vivo* and to build disease models.

## Materials and Methods

### Plasmid constructs

To enable recombinant protein production in insect and mammalian cells using a single set of transfer vectors, we modified the MultiBac expression vector suite ^32,61^ by replacing existing promoters with a dual CMV-p10 promoter ^33^ to derive the acceptor vector pAMI, and the three donor vectors pMIDC, pMIDK, and pMIDS (Extended Data Fig. 1). Open reading frames encoding the TSEN subunits TSEN2 (UniProtKB Q8NCE0), TSEN15 (UniProtKB Q8WW01), TSEN34 (UniProtKB Q9BSV6), TSEN54 (UniProtKB Q7Z6J9), and CLP1 (UniProtKB Q92989) were amplified by polymerase chain reaction (PCR) and cloned into the modified MultiBac vectors leading to pAMI_CLP1, pAMI_TSEN2, pMIDC_TSEN54, pMIDK_TSEN15, and pMIDS_TSEN34. An N-terminal His_6_-tag followed by a Tobacco Etch Virus (TEV) protease cleavage site was engineered in vectors encoding CLP1, TSEN2, and TSEN15, leading to pAMI_His_6_-TEV-CLP1, pAMI_His_6_-TEV-TSEN2, and pMIDK_His_6_-TEV-TSEN15, respectively. Furthermore, a pMIDK plasmid encoding TSEN15 with an N-terminal TEV protease-cleavable Strepavidin-binding peptide (SBP) tag was generated (pMIDK_SBP-TEV-TSEN15). The PCH-causing mutations Tyr^309^Cys (TSEN2), His^116^Tyr (TSEN15), Arg^58^Trp (TSEN34), Ser^93^Pro (TSEN54), and Ala^307^Ser (TSEN54), and the active site mutations His^255^Ala (TSEN34), and His^377^Ala (TSEN2) were introduced via QuikChange mutagenesis. For crystallographic purposes, the coding sequences of TSEN34 (residues 208-310) and TSEN15 (residues 23-170) were cloned into pAMI and pMIDK, respectively, attaching an N-terminal His_10_-tag followed by a TEV protease cleavage site to TSEN15. Prior to integration into the EMBacY baculoviral genome via Tn7 transposition ^61^, acceptor and donor vectors were concatenated by Cre-mediated recombination utilizing the LoxP sites present on each vector. For co-expression of the TSEN15–34 heterodimer, the vectors pMIDK_His_6_-TEV-TSEN15 and pMIDS_TSEN34 were concatenated with the vector pADummy, which was generated by removing the CMV-p10-SV40 expression cassette from pAMI by cleavage with *Avr*II and *Spe*I restriction enzymes and re-ligation of the backbone.

For two-color pre-tRNA cleavage assays, TSEN/CLP1-FLAG and TSEN-STREP wt complexes were cloned into pBIG2ab and pBIG1a expression vectors, respectively, using the biGBac cloning system ^62^. TSEN2^H377A^ and TSEN34^H255A^ point mutants were generated using the Q5 site-directed mutagenesis kit (New England Biolabs) prior to assembly into biGBac vectors, generating both the TSEN/CLP1-FLAG (TSEN2^H377A^) and TSEN/CLP1-FLAG (TSEN34^H255A^) pBIG2ab constructs.

Yeast and human pre-tRNA genes were amplified by PCR from genomic DNA of *Saccharomyces cerevisiae* strain S288C and human embryonic kidney (HEK293) cells, respectively. Pre-tRNA sequences were optimized for *in vitro* transcription (GG at the starting position, CC at pairing position in acceptor stem) and flanked by a preceding T7 promoter sequence and a *Bst*NI cleavage site at the 3′ end of each pre-tRNA. DNA fragments were cloned into a pUC19 vector via sticky end ligation using *Bam*HI and *Hind*III restriction sites. Mature tRNA sequences were obtained by deleting the intron sequence using the Q5 Site-Directed Mutagenesis kit (New England Biolabs). All constructs in this study were verified by Sanger sequencing.

### Production and purification of human TSEN complexes

Recombinant human TSEN complexes were overexpressed in *Spodoptera frugiperda* (*Sf*) 21 cells essentially as described ^32,61,63^. In brief, transfer plasmids encoding TSEN subunits were created by Cre-mediated recombination and recombinant baculoviral BACs were generated by Tn7 transposition in *Escherichia coli* DH10EMBacY cells (Geneva Biotech). *Sf*21 cells were grown in Sf-900 II SFM medium (Thermo Fischer Scientific), transfected with recombinant EMBacY BACs using X-tremeGENE DNA Transfection Reagent (Roche), and incubated for 72 h at 28 °C. Recombinant initial baculoviruses (V_0_) were harvested from cell supernatants and used for production of amplified baculovirus (V_1_) in *Sf*21 suspension cultures at a multiplicity of infection (MOI) < 1. Typically, TSEN complexes were produced in 1.6 liters of *Sf*21 suspension culture at a cell density of 1 × 10^6^ cells ml^−1^ by infection with 0.5-1% (v/v) of V_1_ baculovirus supernatant. 72 h post cell proliferation arrest, insect cells were harvested by centrifugation at 800g for 5 min. Cell pellets were flash-frozen in liquid nitrogen and stored at −80 °C until further use.

Insect cell pellets were resuspended in 10 ml of lysis buffer comprising 50 mM HEPES-NaOH, pH 7.4, 400 mM NaCl, 10 mM imidazole, 1 mM phenylmethanesulfonyl fluoride (PMSF), 1 mM benzamidine, per 100 ml expression volume and lysed by sonication. Lysates were cleared by centrifugation at 37,000 rpm for 40 min in a Type 45 Ti fixed-angle rotor (Beckman Coulter). Pre-equilibrated Ni^2+^-nitrilotriacetic acid (NTA) agarose resin (Thermo Fisher Scientific) was added to the soluble fraction and incubated for 45 min at 4 °C under agitation. Agarose resin was recovered by centrifugation and washed extensively in lysis buffer without protease inhibitors. Bound proteins were eluted in 50 mM HEPES-NaOH, pH 7.4, 400 mM NaCl, 250 mM imidazole. Eluates of immobilized metal ion affinity chromatography (IMAC) were diluted to 150 mM NaCl and loaded onto a HiTrap Heparin HP column (GE Healthcare). Protein complexes were eluted by a linear salt gradient from 150 mM to 2 M NaCl. TSEN complexes were subjected to TEV protease cleavage (1:50 protease to protein mass ratio) at 4 °C to remove the His-tag, concentrated by ultrafiltration using Amicon Ultra centrifugal filters (Merck) with a molecular weight cut-off (MWCO) of 30 kDa and polished by size exclusion chromatography on a Superdex 200 Increase 10/300 GL column (GE Healthcare) equilibrated in 50 mM HEPES-NaOH, pH 7.4, 400 mM NaCl. Peak fractions were pooled, concentrated by ultrafiltration, and flash-frozen in liquid nitrogen after supplementation with 15% (v/v) glycerol.

TSEN15–34 was typically purified from 1.6 liters of infected *Sf*21 suspension culture as stated above but leaving out the heparin chromatography step. IMAC eluates were buffer exchanged into 25 mM HEPES-NaOH, pH 7.4, 400 mM NaCl on a PD-10 desalting column (GE Healthcare), supplemented with TEV protease (1:50 protein to protease mass ratio), concentrated by ultrafiltration using Amicon Ultra (10 kDa MWCO) centrifugal filters (Merck) and polished on a Superdex 200 Increase 10/300 GL column (GE Healthcare) in 25 mM HEPES-NaOH, pH 7.4, 500 mM NaCl. Peak fractions were pooled, concentrated at room temperature to 25 mg ml^−1^ by ultrafiltration, and diluted to 250 mM NaCl and a final protein concentration of 12 mg ml^−1^ for crystallization trials.

For two-color pre-tRNA cleavage assays, viral bacmids encoding wt TSEN-STREP, wt TSEN/CLP1- FLAG, TSEN/CLP1-FLAG (TSEN2^H377A^) and TSEN/CLP1-FLAG (TSEN34^H255A^) pBIG2ab constructs were generated using the Tn7 transposition system in DH10EMBacY cells. The resulting bacmids were transfected into *Sf*9 insect cells using cellfectin II (Gibco). Virus was harvested after 3 days and used to further amplify the viral concentration in a larger *Sf*9 cell culture. Following amplification, protein complexes were expressed in High Five cells for 72 hours at 28 °C and 130 rpm which were subsequently harvested by centrifugation at 1,000 *x g*. Cell pellets were resuspended in purification buffer comprising 20 mM HEPES, pH 8.0, 150 mM NaCl, 1 mM MgCl_2_ and lysed using multiple passes through a dounce homogenizer followed by sonication. Lysate was cleared via centrifugation at 28,000 *x g* for 40 min at 4 °C followed by filtration through a 0.45 μm filter. Purification of TSEN/CLP1-FLAG, TSEN/CLP1-FLAG (TSEN2^H377A^) and TSEN/CLP1-FLAG (TSEN34^H255A^) constructs was carried out via FLAG purification, using the FLAG tag carried by the CLP1 subunit. Lysate was incubated with anti-DYKDDDDK G1 affinity beads (Genscript) for 3 hours at 4 °C and washed with 20 column volumes of purification buffer. Recombinant protein was eluted using 20 column columns purification buffer supplemented with 1 μM DYKDDDDK FLAG peptide (Genscript). Affinity purification of the TSEN-STREP construct was carried out using the STREP tag carried by the TSEN2 subunit. Cleared lysate was loaded onto a StrepTrap HP column (GE Healthcare) pre-equilibrated with purification buffer. Protein was eluted using purification buffer supplemented with 5 mM D-desthiobiotin (Sigma). Following affinity purification, protein-containing fractions were pooled and loaded onto a HiTrap Q column. Protein complexes were eluted in a linear gradient from 150 mM to 2 M NaCl in 20 mM HEPES, pH 8.0, 1 mM MgCl_2_. TSEN-containing fractions were pooled and loaded onto a Superose 6 Increase 10/300 GL column (GE Healthcare) pre-equilibrated in purification buffer. Purified TSEN complexes were analyzed by SDS-PAGE and western blotting.

For overproduction of heterotetrameric TSEN-SBP and TSEN-SBP (TSEN15^H116Y^), adherent human embryonic kidney (HEK) 293T cells were transfected with the expression plasmids with branched polyehtyleneimine (PEI, Sigma-Aldrich). In detail, 4 × 10^6^ HEK293T cells were seeded the day before transfection in 100 mm dishes in DMEM medium (Gibco Life Technologies) with 10% fetal bovine serum (FBS, Capricorn Scientific) and incubated at 37 °C, 5% CO_2_ and 90% humidity. After 24 h, cells were transfected with 13 μg of DNA and a 1:4 ratio of PEI per 100 mm dish. Transfected cells were further incubated for 48 h, detached by addition of Trypsin-EDTA (Sigma Aldrich) and harvested by centrifugation at 500 *x g* for 5 min. The cell pellets were flash frozen in liquid nitrogen and stored at −80 °C until further use. Frozen cell pellets were thawed and resuspended in 1 ml of lysis buffer containing 50 mM HEPES-NaOH, pH 7.4, 400 mM NaCl, 0.5 mM PMSF, 1.25 mM benzamidine, per 100 mm dish and lysed by sonication. Lysates were cleared by centrifugation at 20,817 *x g* for 1 h. Pre-equilibrated High Capacity Streptavidin agarose resin (Pierce) was added to the soluble fraction and incubated for 1 h at 4 °C under agitation. Agarose resin was recovered by centrifugation and washed extensively in lysis buffer without protease inhibitors. Bound proteins were eluted in 50 mM HEPES-NaOH, pH 7.4, 400 mM NaCl, 2.5 mM biotin. TSEN complex eluates were subjected to TEV protease cleavage (1:20 protease to protein mass ratio) at 4 °C to remove the SBP-tag and polished by size exclusion chromatography on a Superdex 200 Increase 3.2/300 column (GE Healthcare) equilibrated in 50 mM HEPES-NaOH, pH 7.4, 400 mM NaCl. Peak fractions were pooled and subjected to pre-tRNA cleavage assays and differential scanning fluorimetry.

### Native mass spectrometry

The buffer of purified TSEN complexes (50 μl at 1.09 mg ml^−1^ for wt TSEN and 1.88 mg ml^−1^ for wt TSEN/CLP1) was exchanged for 200 mM ammonium acetate buffer, pH 7.5, using 30 kDa MWCO centrifugal filter devices (Vivaspin, Sartorius). Native MS experiments were performed on a Quadrupole Time-of-flight (Q-ToF) Ultima mass spectrometer modified for transmission of high mass complexes (Waters, Manchester, UK) ^64^. For data acquisition, 3-4 μl of the sample were loaded into gold-coated capillary needles prepared in-house ^65^. Mass spectrometric conditions were capillary voltage, 1.7 kV; cone voltage, 80 V; RF lens voltage, 80 V; collision energy, 20 V; Aperature3, 13.6. Mass spectra were processed using MassLynx 4.1. At least 100 scans were combined. Acquired mass spectra were calibrated externally using 100 mg ml^−1^ cesium iodide solution. At least 100 scans were combined. Mass spectra were processed in MassLynx and analyzed using Massign ^66^.

### Phosphoprotein analysis

To analyze the phosphorylation state of TSEN subunits, 50 μg of purified protein complexes were treated with 2,000 U of Lambda Protein Phosphatase (New England Biolabs) in 200 μl dephosphorylation buffer (50 mM HEPES-NaOH, pH 7.4, 400 mM NaCl, 1 mM DTT, 1 mM MnCl_2_) for 2 h at 30 °C. Untreated and dephosphorylated complexes were analyzed via SDS-PAGE. Gels were stained with ProQ Diamond Phosphoprotein Gel Stain (Thermo Fisher Scientific) according to the manufacturer’s instructions and imaged on a Typhoon Bioimager (GE Healthcare) at excitation and emission wavelengths of 532 nm and 560 nm, respectively. Imaged gels were subsequently stained with InstantBlue Coomassie (Expedeon).

### Nuclear Extracts

To assay pre-tRNA splicing using patient fibroblasts, we prepared nuclear extracts. Cells from at least four confluent 15 cm dishes were trypsinized, the cell pellet washed once with PBS and spun for 2 min at 1,200 rpm. The pellet was re-suspended in 1 ml 1 x PBS and transferred to a 1.5 ml tube. The tubes were centrifuged for 5 min at 1,200 rpm. The pellet was re-suspended in one volume Buffer A (10 mM HEPES-KOH pH 8.0, 1 mM MgCl_2_, 10 mM KCl, 1 mM DTT) and incubated for 15 min on ice. A 1-ml syringe (fitted with a 0.5 mm × 16 mm needle) was filled with Buffer A and thereafter fully displaced by the plunger to remove all the remaining air within the syringe. Cells were lysed by slowly drawing the suspension into the syringe followed by rapidly ejecting against the tube wall. This step was repeated five times for complete lysis to occur. The sample was then spun for 20 s at 13,000 rpm. The pellet was re-suspended in two-thirds of one packed cell volume in Buffer C (20 mM HEPES-KOH, pH 8.0, 1.5 mM MgCl_2_, 25 % (v/v) glycerol, 420 mM NaCl, 0.2 mM EDTA, 0.1 mM PMSF, 1 mM DTT) and incubated on ice with stirring for 30 min. The suspension was centrifuged for 5 min at 12,000 rpm. The supernatant (corresponding to nuclear extracts) was dialyzed for 1 h against 30 mM HEPES-KOH, pH 7.4, 100 mM KCl, 5 mM MgCl_2_, 10 % (v/v) glycerol, 1 mM DTT, 0.1 mM AEBSF using dialysis membranes (Millipore ‘V’ series membrane). Afterwards, protein concentrations were determined (BioRad Bradford reagent), normalized using dialysis buffer and immediately used for enzymatic assays or snap-frozen and stored at −80 °C.

### Pre-tRNA cleavage assays

For non-radioactive assays, pUC19 vectors encoding *S.c.* pre-tRNA^Phe^_GAA_2-2, human pre-tRNA^Tyr^_GTA_8-1, *S.c.* tRNA^Phe^_GAA_2-2 and human tRNA^Tyr^_GTA_8-1 were linearized using *Bst*NI and template DNA was isolated by agarose gel electrophoresis. RNA substrates were produced by run-off *in vitro* transcription using T7 RNA polymerase (New England Biolabs) and purified via anion exchange chromatography as described before with slight modifications ^36,67^. Briefly, 1 μg ml^−1^ of template DNA was mixed with 1000 U ml^−1^ of T7 polymerase and 1.5 mM of each rNTP (New England Biolabs) in 40 mM Tris-HCl, pH 7.9, 9 mM MgCl_2_, 2 mM spermidine, 1 mM DTT, and incubated for 4 h at 37 °C. To isolate transcribed RNAs, the reaction mixture was diluted in a 1:1 ratio (v/v) with AEX buffer comprising 50 mM sodium phosphate, pH 6.5, 0.2 mM EDTA, and loaded onto a HiTrap DEAE FF column (GE Healthcare) equilibrated in AEX buffer and eluted by a linear gradient from 0 to 700 mM NaCl. RNA containing fractions were analyzed via denaturing Urea-PAGE with subsequent toluidine blue staining. RNAs were concentrated by ultrafiltration using Amicon Ultra 3 MWCO centrifugal filters (Merck) and stored at −20 °C. 1 μg TSEN complexes were mixed with the respective RNA in a 1:5 molar ratio in 50 mM HEPES-NaOH, pH 7.4, 100 mM NaCl, 2 mM MgCl_2_, 1 mM DTT in a 20 μl reaction volume and incubated at 37 °C for 45 min. Reactions were stopped by the addition of a 2x RNA loading buffer (95% formamide, 0.02% SDS, 1 mM EDTA) and incubation at 70 °C for 10 min. Reaction products were separated by denaturing Urea-PAGE and visualized by toluidine blue staining.

For pre-tRNA cleavage assays using radioactive probes, *S.c.* pre-tRNA^Phe^_GAA_ (plasmid kindly provided by C. Trotta) was transcribed in vitro using the T7 MEGAshortscript kit (Ambion) including 1.5 MBq [α^32^P]-guanosine-5′-triphosphate (111 TBq/mmol, Hartmann Analytic) per reaction. The pre-tRNA was resolved in a 10% denaturing polyacrylamide gel, visualized by autoradiography and passively eluted from gel slices overnight in 0.3 M NaCl. RNA was precipitated by the addition of three volumes of ethanol and dissolved at 0.1 μM in buffer containing 30 mM HEPES-KOH, pH 7.3, 2 mM MgCl_2_, 100 mM KCl. To assess pre-tRNA splicing, one volume of 0.1 μM body labeled *S. cerevisiae* pre-tRNA^Phe^, pre-heated at 95°C for 60 sec and incubated for 20 min at room temperature, was mixed with four volumes of reaction buffer (100 mM KCl, 5.75 mM MgCl_2_, 2.5 mM DTT, 5 mM ATP, 6.1 mM Spermidine-HCl pH 8.0 (Sigma), 100 U ml^−1^ RNasin RNase inhibitor (Promega)). Equal volumes of this reaction mixture and cell extracts with a total protein concentration of 6 mg ml^−1^ were mixed and incubated at 30°C. At given time points, 5 μl of the mix were deproteinized with proteinase K, followed by phenol/chloroform extraction and ethanol precipitation. Reaction products were separated on a 10% denaturing Urea-polyacrylamide gel, and tRNA exon formation was monitored by phosphorimaging. Quantification of band intensities was performed using ImageQuant software.

For two-colored pre-tRNA cleavage assays, 5′-cyanine5 (Cy5) and 3′-Fluorescin (FITC) labelled human pre-tRNA^Tyr^_GTA_ 3-1 was purchased from Dharmacon. Pre-tRNA was resuspended in nuclease-free water (New England Biolabs) at 100 μM stock concentration. Prior to use, pre-tRNA stock was diluted 1 in 2 into RNA loading buffer (New England Biolabs) and separated on a 10% acrylamide urea-TBE denaturing gel, with the band corresponding to pre-tRNA excised. The excised bands were crushed using a pipette tip in a 1.5 ml Eppendorf tube and incubated in 300 μl of 20 mM Tris-HCl, pH 8.0, 250 mM KCl, overnight at room temperature. Gel fragments were removed by centrifugation at 17,000 *x g.* Supernatant was transferred to a fresh Eppendorf tube and tRNA precipitated through addition of 4 μl RNA-grade glycogen (Thermo Fisher Scientific) and 1 ml of 100% isopropanol. Precipitate was collected through centrifugation at 17,000 *x g* and the pellet washed in 75% ethanol. The resulting pellet was resuspended in nuclease free water (New England Biolabs) and RNA quantified through measurement of A_260_ prior to storage at −80 °C. Purified pre-tRNA was diluted 1:10 in cleavage buffer (20 mM HEPES, pH 8.0, 100 mM KCl, 2.5 mM Dithiothreitol, 5 mM Spermidine-HCl, 5 mM MgCl_2_). Pre-tRNA was incubated at 90 °C for 1 min and cooled to room temperature for 20 min to ensure folding. 20 pmols of folded pre-tRNA substrate was incubated with a final concentration of 5 U ml^−1^ RNasin plus inhibitor (Promega), 5 mM ATP and 8 pmols of TSEN complex in a final reaction volume of 20 μl for 1 h at 30 °C. RNA was extracted through addition of 150 μl of cleavage buffer followed by 150 μl of 25:24:1 phenol:chloroform:isoamyl alcohol solution (Thermo Fisher Scientific). Samples were mixed and centrifuged at 17,000 *x g* for separation of RNA and protein layers. The top layer was transferred to a fresh Eppendorf tube and RNA precipitated through addition of 4 μl RNA-grade glycogen (Thermo Fisher Scientific) and 1 ml of 100% isopropanol. Precipitated RNA was centrifuged at 17,000 *x g* and the pellet washed in 75% ethanol solution. RNA was resuspended in 10 μl of nuclease free water (New England Biolabs). 5 μl of RNA solution was suspended in 5 μl of RNA loading buffer (95% (v/v) formamide, 10 mM EDTA). Samples were boiled at 95 °C for 10 min prior to loading on a 10% acrylamide urea-TBE denaturing gel. Results were visualized using a Typhoon FLA 9000 (GE Healthcare).

### Fluorescent 3′ end labeling of RNA

RNAs were labeled site-specifically at their 3′ ends using periodate chemistry and a hydrazide derivate of cyanine5 (Cy5) fluorophore (Lumiprobe) as described previously ^68^. Typically, 5 μM of RNA were mixed with 2.5 μl of 400 mM NaIO_4_, 13.33 μl of 3 M KOAc, pH 5.2, in a total volume of 400 μl and incubated for 50 min on ice to oxidize the 2′-3′ diols of the terminal ribose to aldehydes. Oxidized RNAs were ethanol precipitated and resuspended in 400 μl of diethylpyrocarbonate (DEPC)-treated water containing 1 mM of Cy5-hydrazide and 13.33 μl of 3 M KOAc, pH 5.2. After incubation at 4 °C overnight in the dark under agitation, RNA was ethanol precipitated and buffer exchanged to fresh DEPC-treated water using a Zeba Spin desalting column (Thermo Fisher Scientific) to remove the unreacted dye. The optical density at wavelengths of 260 nm and 650 nm was measured using a NanoDrop 1000 spectrophotometer (Thermo Fisher Scientific) to determine the frequency of incorporation (FOI; the number of incorporated fluorophores per 1000 nucleotides) and labeling efficiency.

### tRNA pull-down assays

25 μl of the monoclonal α-His antibody (Catalog-ID H1029, Sigma-Aldrich) were mixed with 100 μl buffer comprising 50 mM HEPES-NaOH, pH 7.4, 400 mM NaCl (HS buffer) and coupled to 25 μl of Protein G Agarose (Thermo Fischer Scientific) for 30 min at 4 °C under agitation. Beads were washed twice with 1 ml HS buffer (1500x g, 3 min) and incubated with 8 μg of inactive His-tagged tetrameric TSEN complex (His_6_-tag on TSEN15) in a total volume of 100 μl for 1h at 4 °C. After washing three times with 150 μl buffer comprising 50 mM HEPES-NaOH, pH 7.4, 100 mM NaCl (LS buffer), 100 ng of Cy5-labeled RNA were added to the beads and incubated for 1h at 4 °C under agitation. After binding, beads were washed again 3x in 150 μl LS buffer and bound macromolecules were eluted by addition of 5 μl 4x SDS loading buffer plus 20 μl LS buffer and incubation at 70 °C for 3 min. Eluted components were separated by SDS-PAGE and visualized by in-gel fluorescence on an ImageQuant LAS 4000 system and immunoblotting. As positive and negative controls, the pull-down assay was performed without the addition of inactive tetrameric TSEN to the antibody-coupled beads or in the presence of an excessive amount (2 μg) of unlabeled RNA, respectively.

### Electrophoretic mobility shift assays

3′-Cy5-labelled pre-tRNA substrates (10 nM final) were mixed with increasing amounts of inactive tetrameric TSEN complexes (typically 10 nM up to 1 μM) in a total volume of 20 μl EMSA buffer comprising 50 mM HEPES-NaOH, pH 7.4, 100 mM NaCl, 1 mM DTT, 4% (v/v) glycerol in DEPC-treated water. After incubation for 30 min on ice in the dark, samples were loaded onto a 4% Tris-Borate-EDTA native polyacrylamide gel, which had been pre-run for 15 min at 180V in 0.5x TBE buffer. Free and complexed RNAs were separated for 1 h at 180V at 4 °C in the dark. In-gel fluorescence was detected on an ImageQuant LAS4000 or Typhoon 9400 device (GE Healthcare) to visualize labeled RNA.

### Fluorescence anisotropy measurements

Fluorescence anisotropy measurements were conducted on a Fluorolog-3 spectrofluorometer (Horiba) equipped with automated polarization filters at a controlled temperature of 22 °C. 120 μl of Cy5-labeled RNA with a concentration of 70 nM in 50 mM HEPES-NaOH, pH 7.4, 100 mM NaCl were titrated with TSEN complexes (1.5 μM stock) in a micro fluorescence cuvette. To avoid dilution effects, the titrant solution contained identical concentrations of the labeled RNAs. After each titration step, the solution was mixed carefully and fluorescence anisotropy was continuously assessed in 15 s increments over a period of 450 s. Anisotropy values of each data point were averaged, plotted in dependency of the protein concentration, and dissociation constants (*K*_D_) were obtained by non-linear curve fitting according to a quadratic equation in Prism 5 (GraphPad Software) to compensate for non-negligible receptor concentrations. Experiments were performed in at least biological duplicates.

### Differential scanning fluorimetry

TSEN complexes were mixed to a final concentration of 1 or 3 μM with 4x SYPRO Orange (Merck) stock in 50 mM HEPES-NaOH, pH 7.4, 400 mM NaCl. Protein unfolding was assessed on a PikoReal96 thermocycler (Thermo Fisher) by measuring SYPRO Orange fluorescence over a temperature gradient from 20 – 95 °C (temperature increment 0.2 °C, hold time 10 s) in a 96-well plate format. Values of technical triplicates were averaged, blank corrected, and apparent unfolding temperatures were determined as the half maximum of a sigmoidal Boltzmann fit in Prism 8 (GraphPad Software). Unfolding temperatures of PCH mutants were compared to wt TSEN complex in technical triplicates to assess their impact on stability and are representative of biological duplicates.

### Size exclusion chromatography multi-angle light scattering

Multi-angle light scattering coupled with size exclusion chromatography (SEC-MALS) was done using a Superdex200 Increase 10/300 GL column (GE Healthcare) at a flow rate of 0.5 ml min^−1^ on an HPLC system composed of PU-2080 pumps, PU-2075 UV detector and degaser (JASCO) connected to a 3-angle miniDAWN TREOS light scattering detector (Wyatt Technology Corporation) and an Optilab T-rEX refractive index detector (Wyatt Technology Corporation). A BSA sample (400 μg) for calibration and 330 μg of TSEN15–34 complex at a concentration of 1.65 mg ml^−1^ were run on a pre-equilibrated column in 25 mM HEPES-NaOH, pH 7.5, 250 mM NaCl filtered through a 0.1 μm pore size VVLP filter (Millipore). The refractive index increment (*dn/dc*) of the TSEN15–34 complex was predicted to 0.188 ml g^−1^ using its amino acid composition ^69^. The extinction coefficient of the TSEN15–34 complex at 280 nm was calculated using the ProtParam server (https://web.expasy.org). Data analysis was accomplished using the ASTRA software package (Wyatt Technology Corporation) across individual peaks using the Zimm’s model for data fitting ^70^.

### Limited proteolysis

Purified, full-length TSEN15–34 complex (0.9 mg ml^−1^) was incubated with trypsin (15 μg ml^−1^) in 50 mM HEPES-NaOH, pH 7.4, 400 mM NaCl for 1h at room temperature. The reaction was stopped by the addition of 1 mM PMSF and the proteolyzed complex was applied to a Superdex 200 Increase 10/300 GL column (GE Healthcare) equilibrated in 50 mM HEPES-NaOH, pH 7.4, 400 mM NaCl. Peak fractions were run out on denaturing 11% SDS-PAGE and visualized by staining with InstantBlue Coomassie (Expedeon).

### Denaturating mass spectrometry

The buffer of TSEN15–34 complexes (10 μl at 1.05 mg ml^−1^ in 10 mM HEPES, pH 7.4, 400 mM NaCl, 0.3x Protease Inhibitor) derived from limited proteolysis was exchanged for 200 mM ammonium acetate, pH 7.5, using 3 kDa MWCO Amicon centrifugal filters (Merck Millipore). For protein denaturation, isopropanol was added to a final concentration of 1% (v/v). Subsequently, the sample was analyzed by direct infusion on a Q Exactive Plus Hybrid Quadrupole-Orbitrap mass spectrometer (Thermo Fisher Scientific) equipped with a Nanospray Flex ion source (Thermo Fisher Scientific). For this, 2–3 μl were loaded into gold-coated capillary needles prepared in-house. MS spectra were recorded in positive ion mode using the following settings: capillary voltage, 2 kV; capillary temperature, 250 °C; resolution, 70.000; S-lens RF level, 50; max injection time, 50 ms; automated gain control, 1·10^6^; MS scan range 1000 – 6000 *m/z*. Approximately 300 scans were combined and the peaks were assigned manually.

### Identification of proteins and protein fragments

Gel electrophoresis was performed using 4-12% NuPAGE Bis-Tris gels according to manufacturer’s protocols (NuPAGE system, Thermo Fisher Scientific). Protein gel bands were excised, and the proteins were hydrolyzed as described previously ^71^. Briefly, proteins were reduced with 10 mM dithiothreitol, alkylated with 55 mM iodoacetamide, and hydrolyzed with Trypsin (Roche). Extracted peptides were dissolved in 2% (v/v) acetonitrile, 0.1% (v/v) formic acid and separated using a DionexUltiMate 3000 RSLCnano System (Thermo Fisher Scientific). For this, the peptides were first loaded onto a reversed-phase C18 pre-column (μ-Precolumn C18 PepMap 100, C18, 300 μm I.D., particle size 5 μm pore size; Thermo Fisher Scientific). 0.1% formic acid (v/v) was used as mobile phase A and 80% (v/v) acetonitrile, 0.1% (v/v) formic acid, as mobile phase B. The peptides were then separated on a reversed-phase C18 analytical column (HPLC column Acclaim® PepMap 100, 75 μm I.D., 50 cm, 3 μm pore size; Thermo Fisher Scientific) with a gradient of 4 - 90% B over 70 min at a flow rate of 300 nl min^−1^. Peptides were directly eluted into a Q Exactive Plus Hybrid Quadrupole-Orbitrap mass spectrometer (Thermo Fisher Scientific). Data acquisition was performed in data-dependent and positive ion modes. Mass spectrometric conditions were: capillary voltage, 2.8 kV; capillary temperature, 275 °C; normalized collision energy, 30%; MS scan range in the Orbitrap, *m/z* 350–1600; MS resolution, 70,000; automatic gain control (AGC) target, 3e6. The 20 most intense peaks were selected for fragmentation in the HCD cell at an AGC target of 1e5. MS/MS resolution, 17,500. Previously selected ions were dynamically excluded for 30 s and singly charged ions and ions with unrecognized charge states were also excluded. Internal calibration of the Orbitrap was performed using the lock mass *m/z* 445.120025 ^72^. Obtained raw data were converted to .mgf files and were searched against the SwissProt database using the Mascot search engine 2.5.1 (Matrix Science).

### Crystallization, structure determination, and validation of a minimal TSEN15-34 complex

Crystals of truncated TSEN15–34 complex (TSEN15 residues 23-170 and TSEN34 residues 208-310) were refined manually at 18°C by mixing equal volumes of protein solution containing 12–15 mg ml^−1^ TSEN15–34 in 25 mM HEPES-NaOH, pH 7.4, 250 mM NaCl, and crystallization solution containing 0.1 M Imidazole/MES, pH 6.5, 20% PEG3350, and 0.2 M MgCl_2_ in a vapor diffusion setup. Crystals were cryoprotected by adding 20% (v/v) glycerol to the reservoir solution and flash-frozen in liquid nitrogen. Diffraction data were collected at 100 K to a resolution of 2.1 Å on beamline P14 of the Deutsches Elektronen-Synchrotron (DESY) and were processed and scaled using the X-ray Detector Software (XDS) package ^73^. Crystals belong to the monoclinic space group P2_1_ with two complexes in the asymmetric unit. The structure of TSEN15-34 was solved by molecular replacement with Phaser ^74^ within the Phenix software package ^75^ using a truncated poly-Ala model of the *Aeropyrum pernix* endonuclease (residues 83-169 of the I chain and residues 93-168 of the J chain) (PDB 3P1Z) ^76^ as a search model. The structures of the two domain-swapped TSEN15-34 dimers were manually built with Coot ^77^ and refined with Phenix ^78^ with good stereochemistry. Statistical quality of the final model was assessed using the program Molprobity ^79^. Structure figures were prepared using PyMOL.

### Cell Culture

Human fibroblasts were cultured at 37 °C, 5% CO_2_ in Dulbecco’s modified Eagle’s medium (Sigma) supplemented with 10% fetal bovine serum (Gibco), 100 U ml^−1^ penicillin and 100 μg ml^−1^ streptoMycin sulfate (Lonza). Cells were split and/or harvested at 80-90% confluency using 0.05% Trypsin–EDTA.

### Northern blotting

Isolation of total RNA from cell lines was performed using the Trizol Reagent (Invitrogen) according to the manufacturer’s instructions. Typically, 4-5 μg of RNA was separated in a 10% denaturing Urea-polyacrylamide gel (20 × 25 cm; Sequagel, National Diagnostics). The RNA was blotted on Hybond-N+ membranes (GE Healthcare) and fixed by ultraviolet cross-linking. Membranes were pre-hybridized in 5x SSC, 20 mM Na_2_HPO_4_, pH 7.2, 7% SDS, and 0.1 mg ml^−1^ sonicated salmon sperm DNA (Stratagene) for 1h at 80 °C (for DNA/LNA probes) or 50 °C (for DNA probes). Hybridization was performed in the same buffer overnight at 80 °C (for DNA/LNA probes) or 50 °C (for DNA probes) including 100 pmol of the following [5’-^32^P]-labeled DNA/LNA probe (Exiqon, Denmark; LNA nucleotides are indicated by “*X”): tRNA^Ile^_TAT_1-1 5’ exon probe, 5’-TA*T AA*G TA*C CG*C GC*G CT*A AC-3’, or the following DNA probe: tRNA^Ile^_TAT_1-1 intron probe, 5’-TGC TCC GCT CGC ACT GTC A-3’. Blots were subsequently washed twice at 50°C with 5x SSC, 5% SDS and once with 1x SSC, 1% SDS and analyzed by phosphorimaging. Membranes were re-hybridized at 50 °C using a DNA probe (5’-GCA GGG GCC ATG CTA ATC TTC TCT GTA TCG-3’) complementary to U6 snRNA to check for equal loading.

### Immunoprecipitation of TSEN components

Antibodies against TSEN2, TSEN34, and TSEN54 ^35^ were affinity-purified, and cross-linked to agarose beads, as described ^80^. Briefly, bead-bound antibodies were incubated in 20 mM dimethylphenol (DMP), 200 mM sodium tetraborate at RT and the reaction was then stopped by transferring the beads to 200 mM Tris-HCl, pH 8.0. After washing 3x with TBS/0.04% Triton-X-100, beads were stored at 4 °C. For immunoprecipitation (IP), total cell lysates were prepared from fresh or frozen cell pellets of primary fibroblasts as described ^81^ Upon centrifugation at 16,000 *x g*, clear lysates were collected, protein concentration was measured, and equal amounts of total protein for each sample were used for the IPs. Upon incubation with cell lysates for 90 min at 4 °C while rotating, TSEN complex-bound beads were washed as described ^80^ and split into two aliquots; one was used for a pre-tRNA splicing assay and the other was boiled in SDS-PAGE loading buffer. Pre-tRNA splicing assay was performed as described above, omitting the proteinase K treatment and the phenol/chloroform extraction and ethanol precipitation steps. Instead, aliquots were collected at indicated time-points in tubes already containing an equal amount of 2 x loading buffer and stored at −20 °C. Protein samples were analyzed by SDS-PAGE and immunoblotting.

### Hydro-tRNA sequencing

tRNA sequencing was performed using the hydro-tRNAseq protocol, as described previously ^4^. Briefly, total RNA from human derived fibroblasts was resolved on 12% urea-polyacrylamide gel, followed by recovery of the tRNA fraction within a size window of 60-100 nt. The eluted fraction was subjected to limited alkaline hydrolysis in 10 mM Na_2_CO_3_ and NaHCO_3_ at 60 °C for 1 hr. The hydrolyzed RNA was dephosphorylated and rephosphorylated to reconstitute termini amenable for sequential adapter ligation. Fragments of 19-35 nt were converted into barcoded cDNA libraries, as described previously ^82^, and sequenced on an Illumina HiSeq 2500 instrument. Adapters were trimmed using cutadapt (http://journal.embnet.org/index.php/embnetjournal/article/view/200/458). Sequence read alignments and analysis was performed as described previously ^4^. Split read counts were used for multimapping tRNA reads. Precursor tRNA reads spanned the junctions between mature sequences and leaders, trailers, or introns.

### Sequence alignments

Sequence alignments were done with Clustal Omega ^83^ and visualized using ESPript 3.0 ^84^. Alignments of pre-tRNAs and tRNAs were manually edited in Jalview ^85^.

### Statistical analysis

Student’s two-tailed nonpaired t tests were carried out to determine the statistical significance of differences between samples. A p value less than 0.05 was considered nominally statistically significant for all tests.

### Patient recruitment and ascertainment

Patients suspected for PCH were submitted to the pediatric neurology of the Academic Medical Centre (AMC) for diagnostics. Primary fibroblast cell lines were generated from skin biopsies taken for diagnostic procedures. As soon as DNA diagnostics became available, patient DNA was subjected to genetic analyses. DNA sequencing confirmed the diagnosis and the mutations were confirmed in the fibroblast lines. All procedures were performed with full consent of the legal representative and approval of the Institutional Review Board (IRB).

## Data availability

Atomic coordinates and structure factors were deposited to the Protein Data Bank (http://www.rcsb.org) under accession number PDB ID 6Z9U. The mass spectrometry proteomics data were deposited to the ProteomeXchange Consortium via the PRIDE partner repository with the dataset identifier PXD019034. Hydro-tRNAseq data were deposited with the Gene Expression Omnibus (GEO) repository under accession code GSE151236. Source data for Figs. 1-6 and Extended Data Figs. 1-6 are provided with the paper.

## Acknowledgments

We thank Rupert Abele for the analysis of SEC-MALS experiments, Jan Erik Schliep for DSF data analysis with the ProteoPlex algorithm, and Imre Berger for providing the MultiBac reagents. The synchrotron MX data were collected at beamline P14 operated by EMBL Hamburg at the PETRA III storage ring (DESY, Hamburg, Germany). We thank Gleb Bourenkov for the assistance in using the beamline. S.T. acknowledges Robert Tampé and all members of his group for discussions and comments on the manuscript and excellent administrative and technical support. S.P. thanks Kristina Uzunova for sharing her expertise in antibody purification and protein biochemistry. M.B. and C.S. acknowledge funding from the Federal Ministry for Education and Research (BMBF, ZIK program, 03Z22HN22), the European Regional Development Funds (EFRE, ZS/2016/04/78115) and the MLU Halle-Wittenberg. This study was furthermore supported by grants of the German Research Foundation (grant number TR 1711/1-7) to S.T., the Austrian Science Fund (grant number FWF P29888) to J.M. and S.T., the CRC 902 Molecular Principles of RNA-based Regulation (S.S. and S.T.), and a Boehringer Ingelheim Fonds fellowship to S.S.

## Author contributions

S.S., P.D., A.P., and S.T. expressed, purified, and prepared protein complexes from insect and mammalian cells. S.S., P.D., A.P., and S.P. performed biochemical assays. E.P.R. cloned and purified FLAG- and STREP-tagged TSEN complexes and performed dual-color pre-tRNA cleavage assays. S.P. and S.W. performed pre-tRNA splicing assays; Northern blots and IP experiments on human fibroblasts were performed by S.P.. M.B. and C.S. conducted MS experiments and analyzed the data. S.S. and S.T. performed crystallography experiments, collected X-ray diffraction data, and built the atomic model. F.B. generated cell lines of PCH patient-derived fibroblast. T.G. performed hydro-tRNAseq experiments and bioinformatic analyses under supervision of T.T.. J.M. and S.T conceived the project, supervised the work, and designed the experiments. S.S., P.D., and S.T. wrote the initial draft of the manuscript with input from all authors. J.M., and S.T. acquired funding.

## Additional information

Correspondence and requests for materials should be addressed to J.M. or S.T.

---

**Extended Data Fig. 1.**
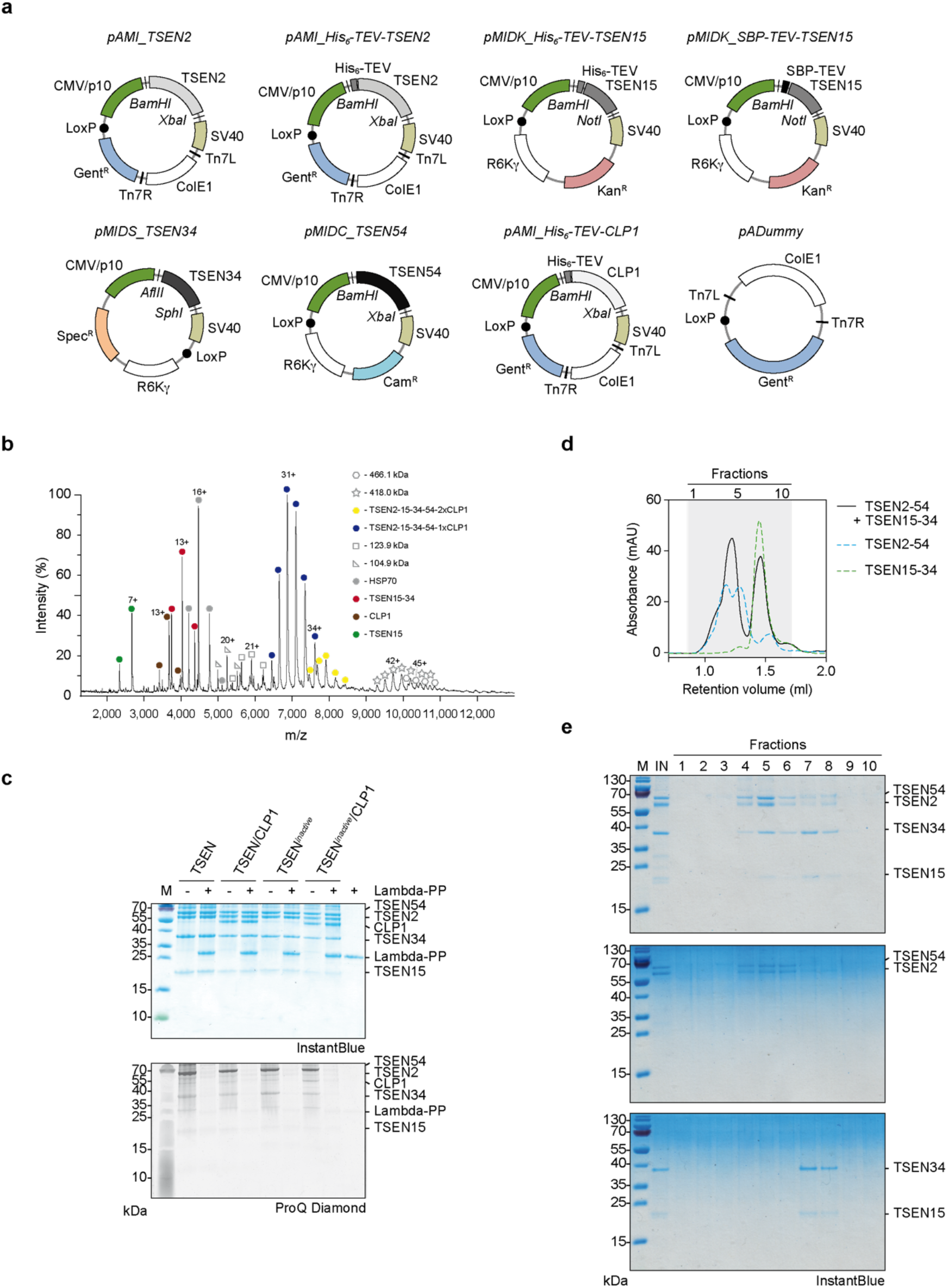
Biochemical characterization of recombinant TSEN and TSEN/CLP1 complexes. **a**, Maps of modified MultiBac vectors encoding TSEN/CLP1 complex components. For expression in mammalian and insect cells, the acceptor vector pAMI and donor vectors pMIDC, pMIDK, and pMIDS carry the CMV/p10 dual promoter. Transcription is terminated by the SV40 poly-A late signal (SV40). The transposase elements Tn7L and Tn7R, the LoxP element (black dot) for Cre-mediated recombination, the origins of replication ColE1 and R6Kγ, and the resistance markers for gentamicin (Gent^R^), chloramphenicol (Cam^R^), kanamycin, (Kan^R^), and spectinomycin (Spec^R^) are shown. Restriction sites, hexahistidine-tags (His_6_), the Streptavidin-binding peptide-tag (SBP), and the TEV cleavage site (TEV) are indicated. **b**, Native mass spectrum of pentameric TSEN/CLP1 complex from an aqueous ammonium acetate solution. Charge states of the predominant TSEN/CLP1 assembly (blue circles), a minor populated TSEN complex with two CLP1 subunits (yellow circles), monomeric CLP1 (brown circles), HSP70 (grey circles), the TSEN15–34 heterodimer (red circles) and TSEN15 (green circles) are indicated. Unidentified protein assemblies are denominated by their molecular weights. **c**, Analysis of phosphorylation states of TSEN components by the phospho-specific ProQ Diamond gel stain. The strong band at 59 kDa in the ProQ Diamond stain corresponds to TSEN54. Lambda-PP, Lambda protein phosphatase. **d**, Assembly assay with TSEN2–54 and TSEN15–34 heterodimers via size exclusion chromatography (SEC). Absorbance profiles (280 nm) of reconstituted TSEN complex (black line), and the heterodimers TSEN2–54 (blue dashed line) and TSEN15–34 (green dashed line) are shown. **e**, SDS-PAGE of SEC fractions (grey area as indicated in **d**) with subsequent InstantBlue staining. Unprocessed gels for **c** and **e** are shown in Source Data 2.

**Extended Data Fig. 2.**
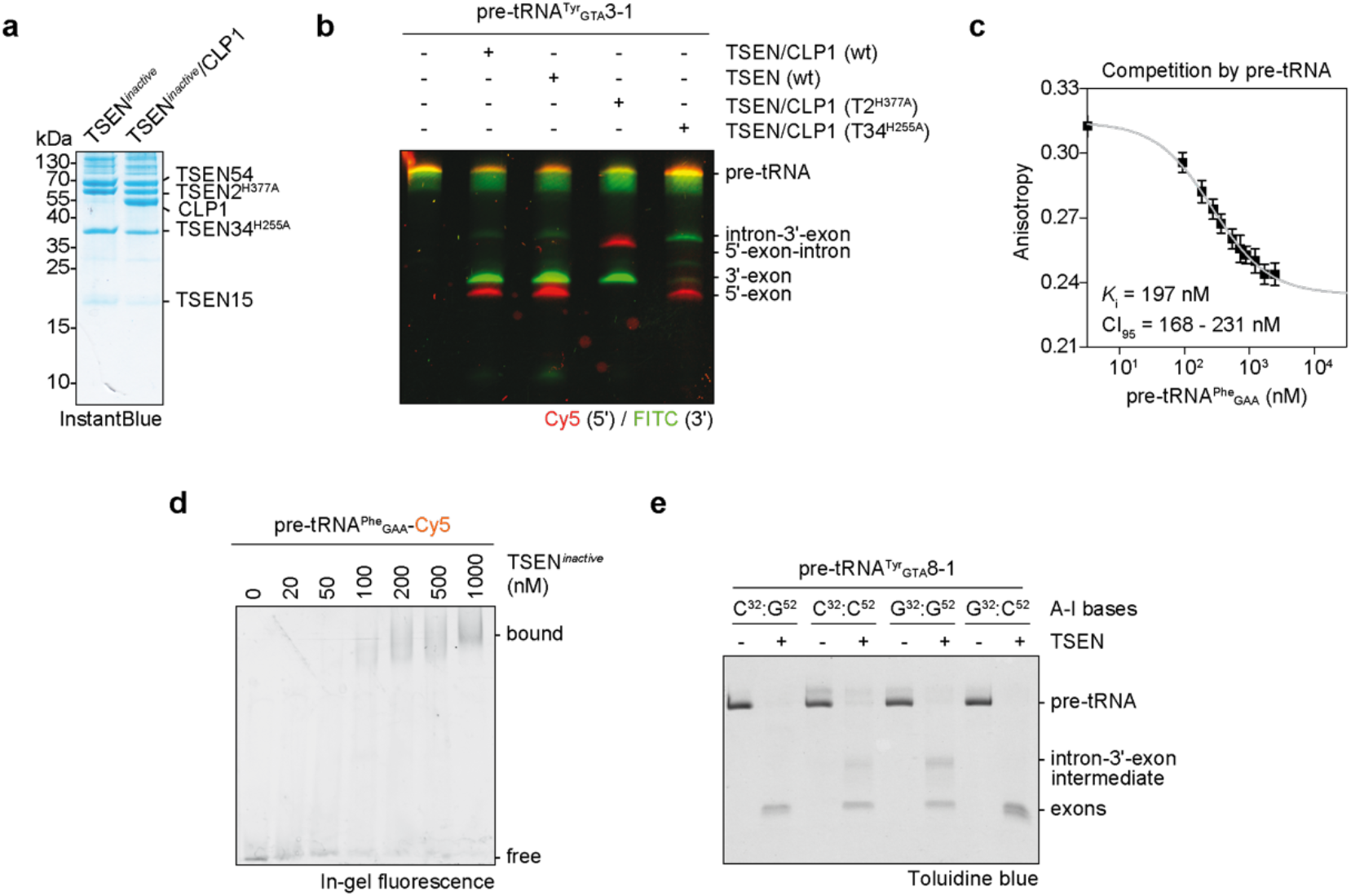
Active involvement of the A–I base pair in coordinating pre-tRNA cleavage. **a**, SDS-PAGE of purified, recombinant inactive TSEN and inactive TSEN/CLP1 complexes (TSEN2^H377A^ and TSEN34^H255A^ double mutant). Protein size markers and protein identities are indicated. **b**, Two-colored pre-tRNA cleavage assay with TSEN-STREP and TSEN/CLP1-FLAG wt complexes and complexes carrying the TSEN2^H377A^ (T2^H377A^) or TSEN34^H255A^ (T34^H255A^) substitution. RNA cleavage products were separated on a denaturing Urea-PAGE and visualized by fluorescence of cyanine5 (Cy5) and Fluorescin (FITC). **c**, Thermodynamic competition parameters deduced from fluorescence anisotropy experiments. Inactive, tetrameric TSEN bound to fluorescently labeled pre-tRNA^Phe^_GAA_ was titrated with unlabeled pre-tRNA. **d**, Electrophoretic mobility shift assay with fluorescently labeled pre-tRNA^Phe^_GAA_ and inactive, tetrameric TSEN (TSEN^*inactive*^). Free and bound fractions of pre-tRNA were analyzed via 4% TBE native PAGE with subsequent in-gel fluorescence measurement. **e**, Impact of A-I base pair mutations in pre-tRNA^Tyr^_GTA_8-1 on endonucleolytic activity by tetrameric TSEN revealed by a pre-tRNA cleavage assay. CI_95_ – 95% confidence interval, C^32^:G^52^ – canonical A-I base pair, C^32^:C^52^ and G^32^:G^52^ – disrupted A-I base pair, G^32^:C^52^ – inverted A-I base pair. Panels are representatives of three independent experiments. Unprocessed gels for **a**, **b**, **d**, and **e** are shown in Source Data 4.

**Extended Data Fig. 3.**
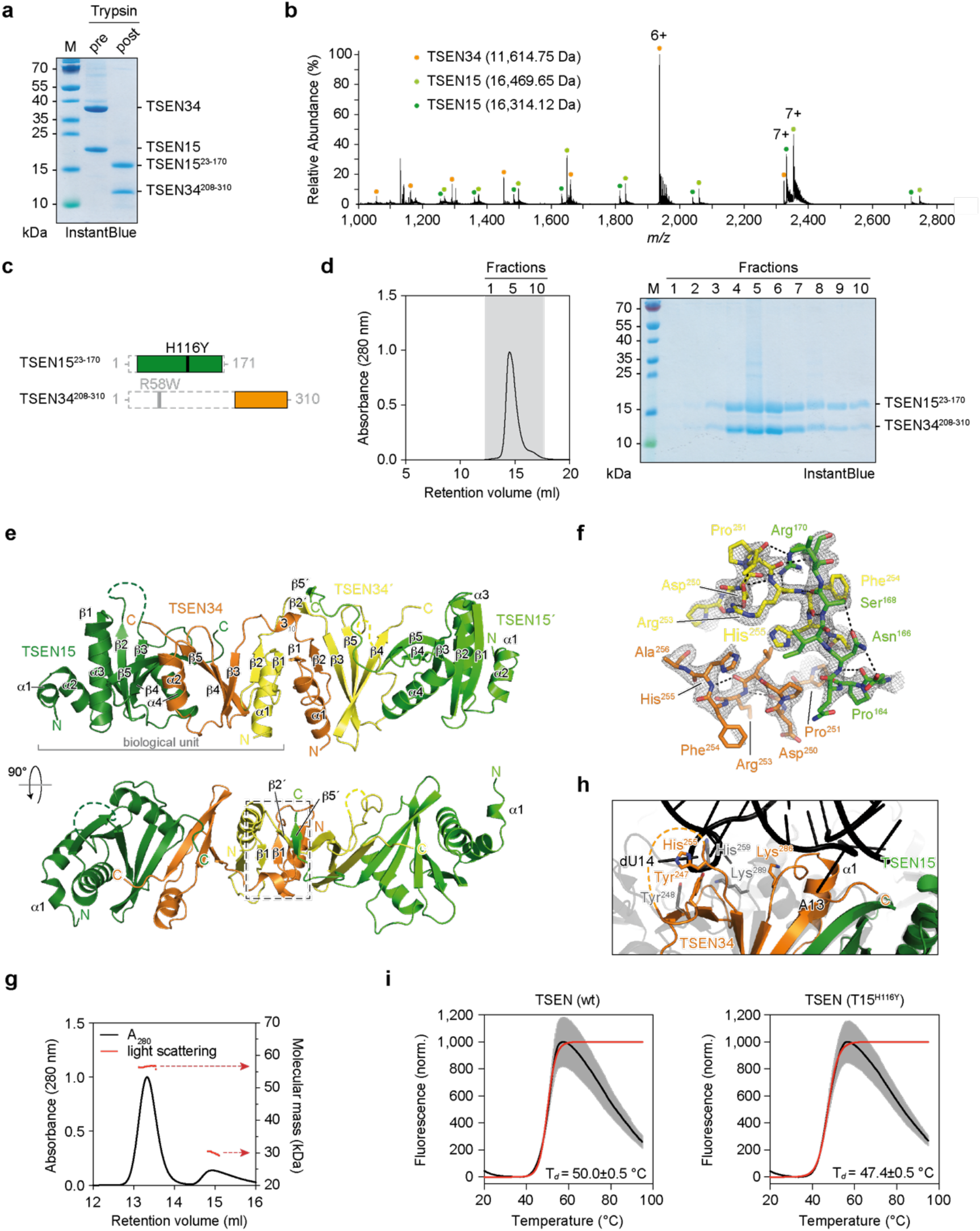
X-ray crystal structure of a TSEN15–34 heterodimer derived by limited proteolysis. **a**, Analysis of the limited tryptic digestion of the TSEN15–34 heterodimer by SDS-PAGE. **b**, Denaturing mass spectrum of the proteolytically stable fragments of the TSEN15–34 heterodimer from an aqueous ammonium acetate solution. The mass spectrum shows the presence of a TSEN34 fragment (orange circles), and two TSEN15 fragments (dark and light green circles) differing in mass only by a C-terminal arginine as revealed by LC-MS/MS. **c**, Bar diagrams of tryptic fragments of TSEN15 and TSEN34. Proteolyzed regions are indicated by dashed boxes. Positions of PCH mutations are shown. **d**, Purification of the re-cloned core of the TSEN15–34 heterodimer via SEC. The absorbance profile at 280 nm is shown. Fractions of the indicated retention range (grey area) were analyzed by SDS-PAGE. **e**, Asymmetric unit of the TSEN15–34 crystal. The biological unit (bracket) and the domain-swap area (dashed box) are indicated. α-helices and β-sheets are numbered for each subunit. **f**, Stick representation of amino acids of the domain-swap area with electron density (2F_o_-F_c_, 1.5σ). **g**, Molecular mass determination by size exclusion chromatography multi-angle light scattering (SEC-MALS) of the TSEN15–34 sample used for crystallization. The data reveal a dominant population of a dimer-of-a-heterodimer (13.2 ml, 56.6 kDa) and a minor populated heterodimer (14.9 ml, 30.0 kDa). Light scattering is shown as red dots. Mass determination by SEC-MALS was confirmed by two independent experiments. **h**, Superposition of the TSEN15–34 heterodimer and the pre-tRNA endonuclease from *Archaeoglobus fulgidus* at the interaction sites with the bulge-helix-bulge RNA (PDB ID 2GJW). Nucleotide positions of the RNA (black) and residues of the catalytic triads are shown. **i**, Representative thermal denaturation curves as shown in Fig. 3g of recombinant wt TSEN and mutant TSEN (T15^H116Y^) complexes derived from DSF. Sigmoidal Boltzmann fits are shown as red lines. Grey zones show standard deviations (SD) from technical triplicates. Denaturation temperature (T_*d*_) is presented with error of fit. Unprocessed gels for **a** and **d** are shown in Source Data 6.

**Extended Data Fig. 4.**
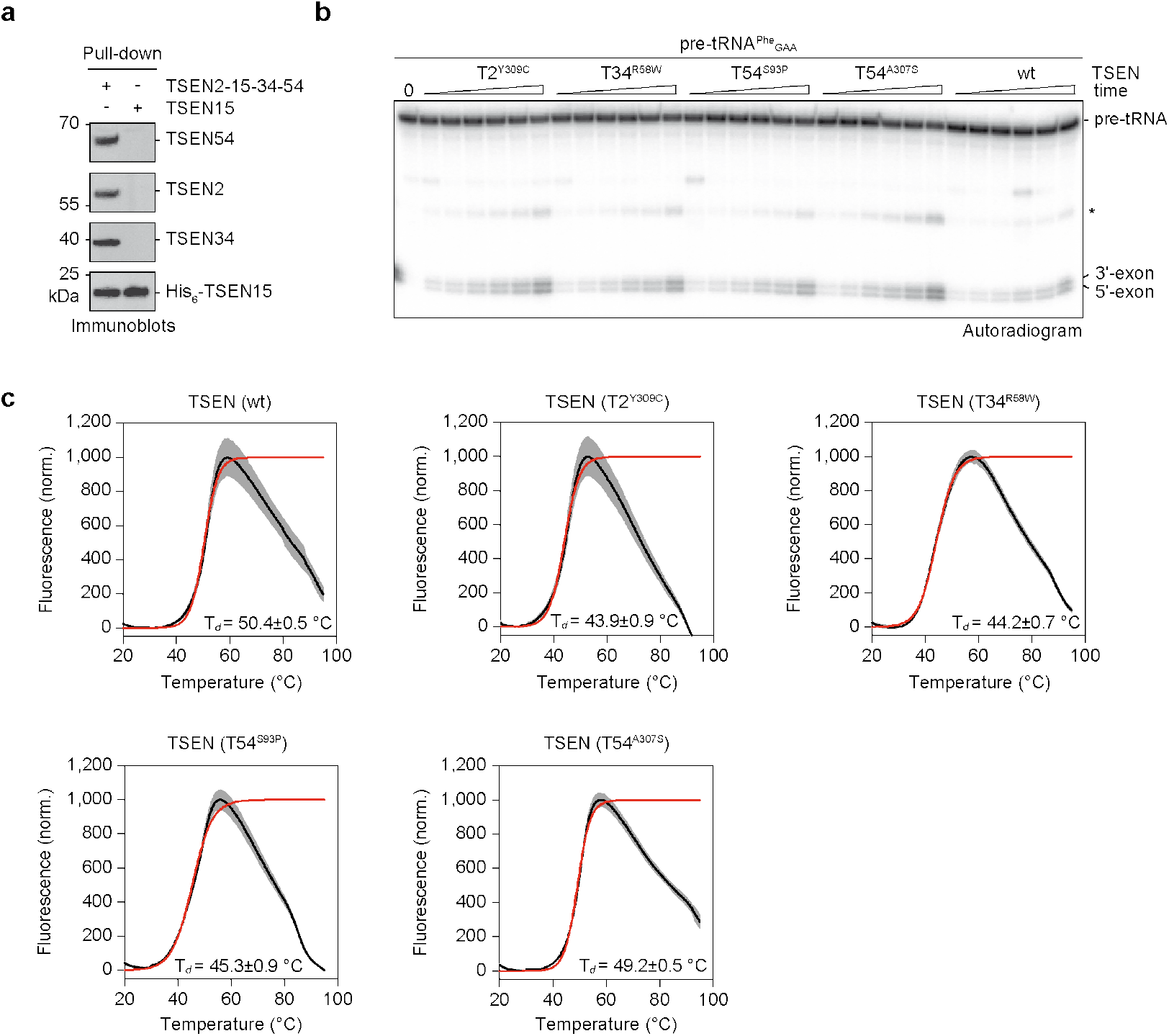
PCH mutations affect thermal stability but not activity of recombinant TSEN *in vitro*. **a**, Pull-down assay from HEK293T cells overexpressing TSEN subunits TSEN2, His_6_-TSEN15, TSEN34, and TSEN54, or His_6_-TSEN15 alone. Co-precipitated proteins were visualized by immunoblotting. **b**, Pre-tRNA cleavage assay (time course) of radioactively labelled *S.c.* pre-tRNA^Phe^_GAA_ with wt or mutant TSEN complexes revealed by phosphorimaging. The asterisk indicates an intermediate cleavage product. **c**, Representative thermal denaturation curves as shown in Fig. 4d of recombinant wt and mutant TSEN complexes derived from DSF with sigmoidal Boltzmann fits as red lines. Grey zones show SDs from technical triplicates. Denaturation temperature (T_*d*_) is presented with error of fit. Unprocessed gels for **a** and **b** are shown in Source Data 9.

**Extended Data Fig. 5.**
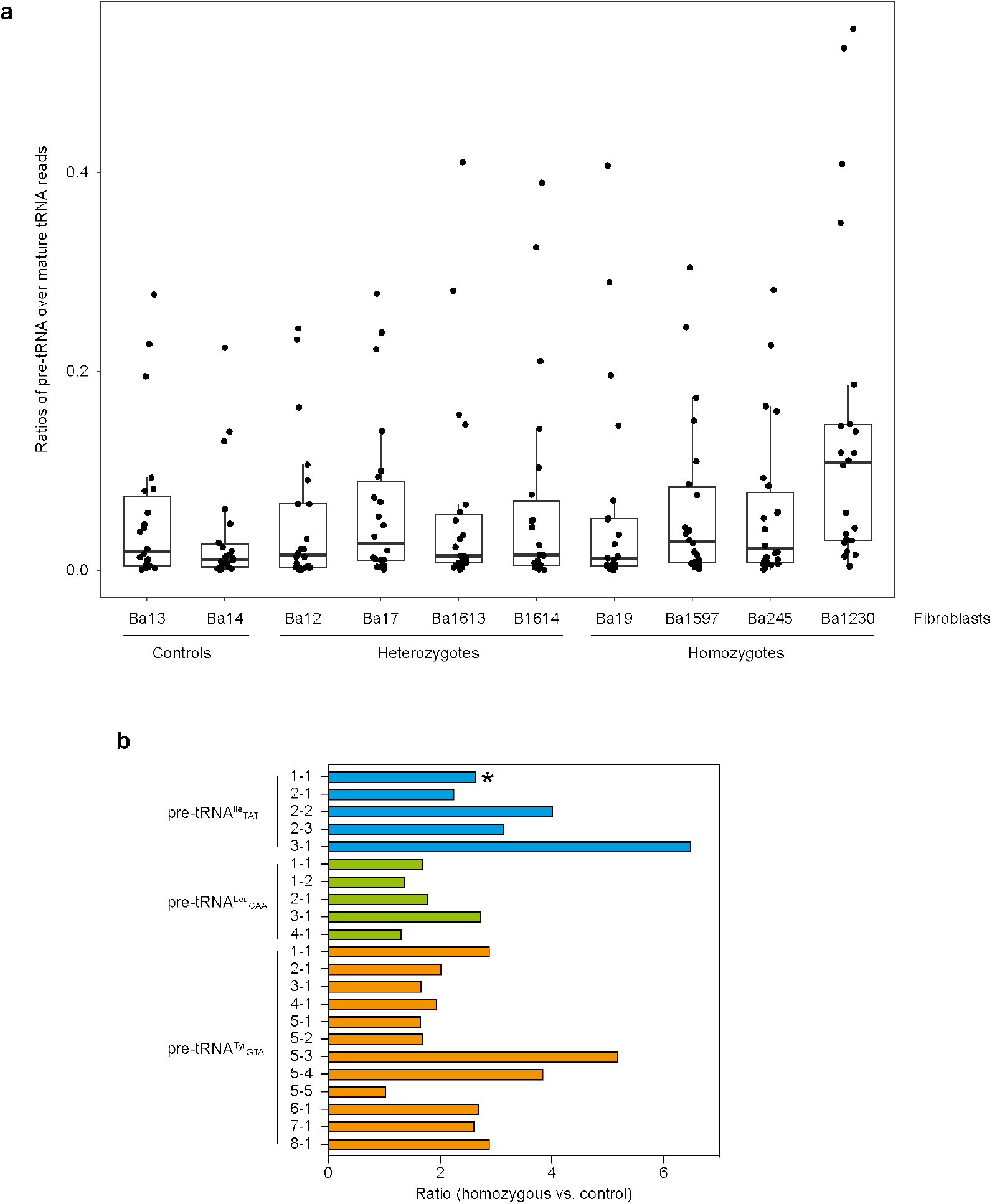
Hydro-tRNAseq reveals accumulation of intron-containing pre-tRNAs in *TSEN54* c.919G>T fibroblasts. **a**, Boxplots showing ratios of pre-tRNA over mature tRNA reads for all intron-containing tRNAs from hydro-tRNAseq of PCH patient-derived and control fibroblasts. Pre-tRNA reads for all samples and tRNAs are less abundant than mature tRNA reads (ratios < 1). The lowest median corresponds to a homozygous control, and the highest median to a homozygous TSEN54^A307S^ patient. **b**, For every intron-containing tRNA, the ratio of hydro-tRNAseq reads mapped to pre-tRNAs over mature tRNAs was calculated. The average ratio of all patients over the average ratio of all homozygous controls is shown. TSEN54 c.919G>T fibroblasts exhibit an increase of pre-tRNA/mature tRNA ratios for all intron-containing tRNAs. tRNA^Ile^_TAT_ isodecoders targeted by a 5’ exon probe shown in Fig. 5b are highlighted in blue. tRNA^Ile^_TAT_1-1 targeted by an intron probe in Fig. 5b,c is marked with an asterisk.

**Extended Data Fig. 6.**
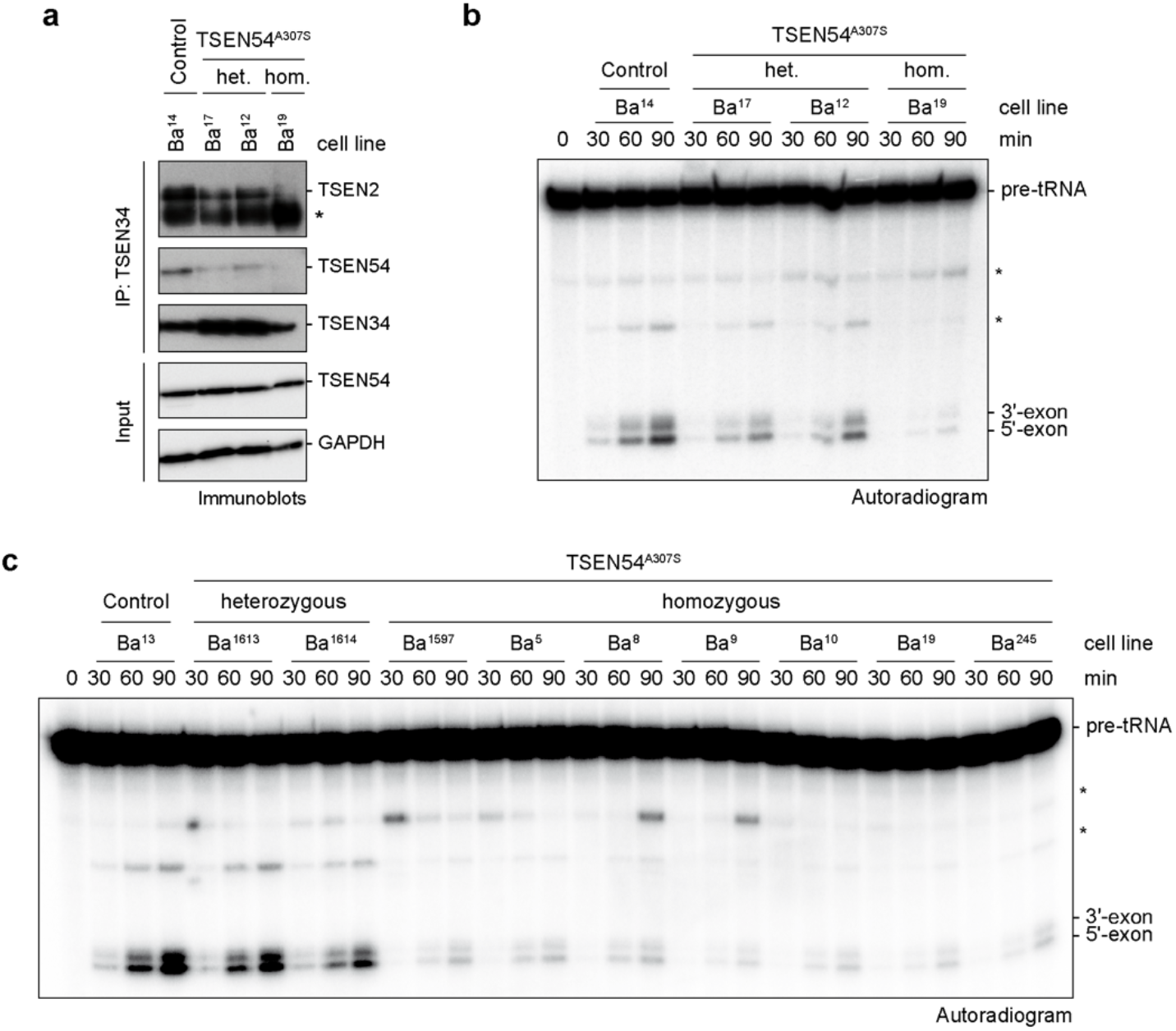
Reduced pre-tRNA cleavage activity in PCH patient-derived cell extracts is associated with altered composition of TSEN. **a**, Co-immunoprecipitation (IP) assay using an α-TSEN34 antibody with cell lysates derived from a control cell line and fibroblast derived from a PCH patient (Ba^19^) and the parents (Ba^17^ and Ba^12^) analyzed by immunoblotting. The asterisk indicates the heavy chain of the α-TSEN34 antibody. GAPDH served as a loading control. **b**, On-bead pre-tRNA cleavage assay (time course) with radioactively labelled *S.c.* pre-tRNA^Phe^_GAA_ and immunoprecipitated TSEN complexes (α-TSEN34 antibody-coupled resin) shown in (**a**). Unspecific bands are indicated by asterisks. **c**, On-bead pre-tRNA cleavage assay (time course) with radioactively labelled *S.c.* pre-tRNA^Phe^_GAA_ and immunoprecipitated TSEN complexes (α-TSEN2 antibody-coupled resin) derived from control fibroblasts and from fibroblasts carrying heterozygous or homozygous *TSEN54 c.919G*>T mutation. Unspecific bands are indicated by asterisks. Data are representative of at least two independent experiments. Unprocessed gels for **a**, **b**, and **c** are shown in Source Data 14 and 15.

