## Supplementary Information for "Assembly defects of the human tRNA splicing endonuclease contribute to impaired pre-tRNA processing in pontocerebellar hypoplasia"

### Supplementary Figures, Tables, and Source Data Files

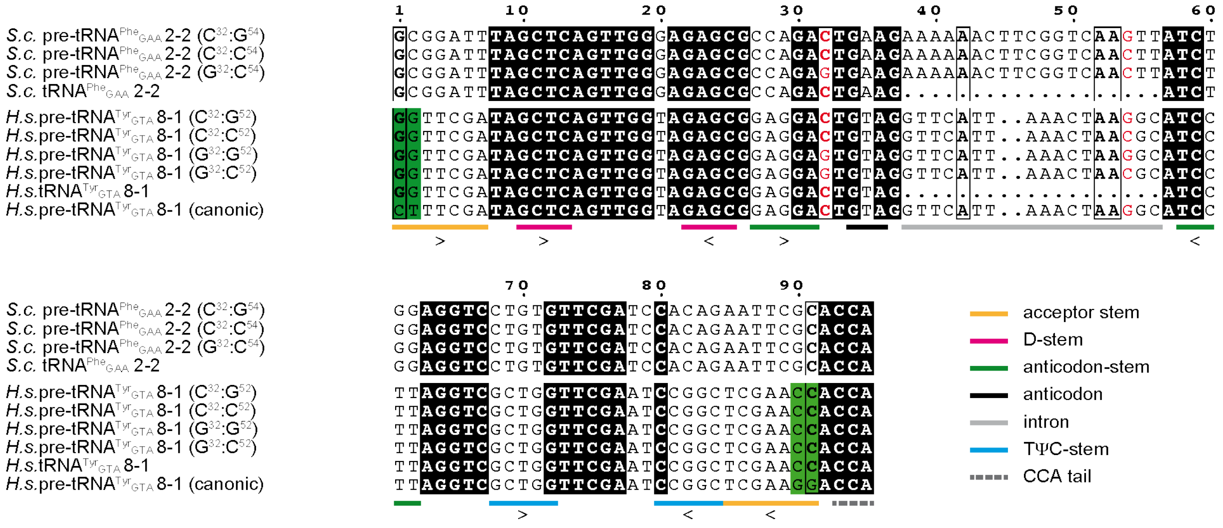

**Supplementary Fig. 1 | Sequence comparison of pre-tRNA and tRNA molecules.** Sequence alignments were performed using Clustal Omega, edited in Jalview and colored by conservation using ESPript 3.0. A–I base pair residues are colored in red. Predicted stem structures, anticodon, intron and CCA tail are indicated by colored bars. Ribonucleotides modified for efficient *in vitro* transcription are boxed in green and compared to the canonical sequence.

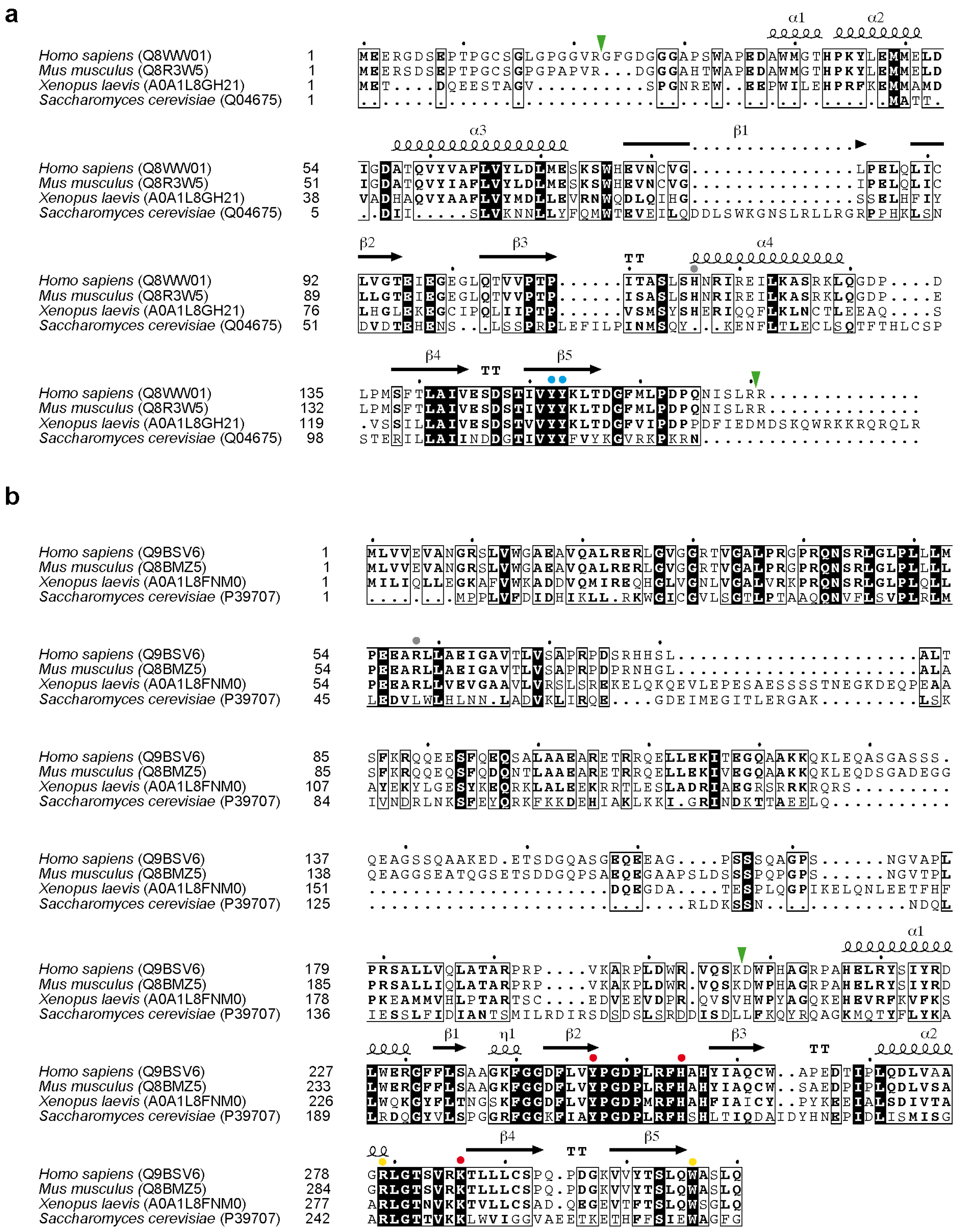

**Supplementary Fig. 2 | Sequence conservation of TSEN15 and TSEN34.** Sequence alignments were performed using Clustal Omega and colored by conservation using ESPript 3.0. **a**, The TSEN15 sequence alignment includes orthologues from *Homo sapiens* (UniProtKB Q8WW01), *Mus musculus* (UniProtKB Q8R3W5), *Xenopus laevis* (UniProtKB A0A1L8GH21), and *Saccharomyces cerevisiae* (UniProtKB Q04675). **b**, The TSEN34 sequence alignment includes orthologues from *Homo sapiens*

(UniProtKB Q9BSV6), *Mus musculus* (UniProtKB Q8BMZ5), *Xenopus laevis* (UniProtKB A0A1L8FNM0), and *Saccharomyces cerevisiae* (UniProtKB P39707). Tryptic sites identified from limited proteolysis experiments are shown by green arrow heads. The YY-motif is indicated by blue circles, residues of the catalytic triad are highlighted by red circles, and residues possibly involved in the cation-$\pi$-interaction are shown as yellow circles. Residues mutated in PCH (TSEN15$^{H116Y}$, TSEN34$^{R58W}$) are indicated by grey circles. Helices and strands are numbered sequentially according to the TSEN15–34 X-ray crystal structure and are indicated above the alignments. TT – $\beta$-turn.

**a**

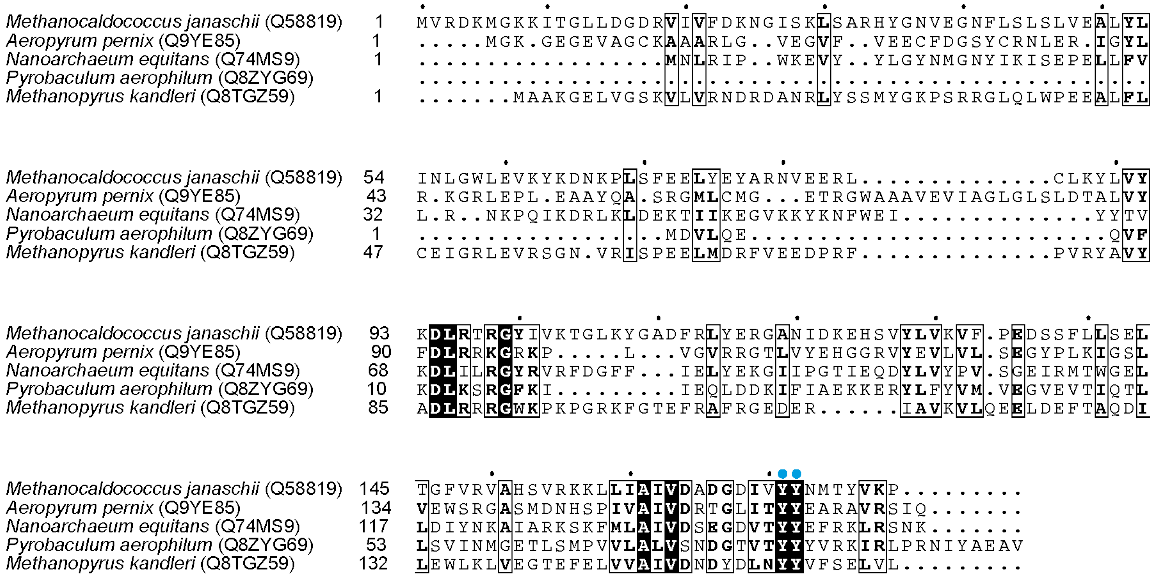

**Supplementary Fig. 3 | Sequence conservation of Archaeal α4 and (αβ)2 endonucleases highlighting the YY-motif.** Sequence alignments were performed using Clustal Omega and colored by conservation using ESPript 3.0. The sequence alignment includes orthologues from *Methanocaldococcus jannaschii* (UniProtKB Q58819), *Aeropyrum pernix* (UniProtKB Q9YE85), *Nanoarchaeum equitans* (UniProtKB Q74MS9), *Pyrobaculum aerophilum* (UniProtKB Q8ZYG6), and *Methanopyrus kandleri* (UniProtKB Q8TGZ5). The YY-motif is indicated by blue dots.

#### Supplementary Tables

**Supplementary Table 1 | Masses of protein subunits and complexes observed in native MS spectra.** The experimentally determined and theoretically calculated masses as well as the mass differences are given. A larger mass difference (*) originates from incomplete desolvation and can be in part attributed to the high phosphorylation state of the TSEN54 subunit (Extended Data Fig. 1c).

| Composition | Experimental mass (Da) | Theoretical mass (Da) | Δ mass (Da) |
|---|---|---|---|
| **TSEN** | | | |
| TSEN2–15–34–54 | 165573 ±130 | 164416 | 1157 (*) |
| *unassigned* | 104865 ±48 | | |
| HSP70 | 71461 ±6 | 71432 | 29 |
| TSEN15–34 | 52389 ±15 | 52350 | 39 |
| TSEN15 | 18693 ±4 | 18698 | -5 |
| **TSEN/CLP1** | | | |
| *unassigned* | 466075 ±127 | | |
| *unassigned* | 417988 ±36 | | |
| TSEN2-15-34-54-2xCLP1 | 261096 ±182 | 259822 | 1274 (*) |
| TSEN2-15-34-54-1xCLP1 | 212967 ±98 | 212119 | 848 (*) |
| *unassigned* | 123948 ±28 | | |
| *unassigned* | 104818 ±6 | | |
| HSP70 | 71521 ±3 | 71432 | 89 |
| TSEN15–34 | 52412 ±7 | 52350 | 62 |
| CLP1 | 47776 ±5 | 47703 | 73 |
| TSEN15 | 18704 ±0 | 18698 | 6 |

| Subunits | Uniprot KB | Theoretical mass (Da) |
|---|---|---|
| TSEN15 | Q8WW01 | 18698 |
| TSEN34 | Q9BSV6 | 33652 |
| TSEN2 | Q8NCE0 | 53247 |
| TSEN54 | Q7Z6J9 | 58819 |
| CLP1 | Q92989 | 47703 |
| HSP70 | Q9U639 | 71432 |

**Supplementary Table 2 | Protein identification by LC-MS/MS.** The protein masses, the number of identified peptide sequences, the number of observed spectra, and the sequence coverage are given for TSEN subunits and CLP1 of purified TSEN, TSEN/CLP1 and proteolyzed TSEN15–34 complexes.

| | | | TSEN | | | TSEN/CLP1 | | |
|---|---|---|---|---|---|---|---|---|
| **Protein** | **UniProtKB** | **Mass** (Da) | **Peptide sequences** (#) | **Spectra** (#) | **Sequence coverage** (%) | **Peptide sequences** (#) | **Spectra** (#) | **Sequence coverage** (%) |
| TSEN15 | Q8WW01 | 18629 | 9 | 140 | 47.4 | 11 | 180 | 73 |
| TSEN34 | Q9BSV6 | 33631 | 53 | 907 | 100.0 | 58 | 1583 | 100 |
| TSEN2 | Q8NCE0 | 53213 | 99 | 1228 | 98.5 | 86 | 1946 | 99 |
| TSEN54 | Q7Z6J9 | 58783 | 67 | 771 | 84.8 | 68 | 1132 | 95 |
| CLP1 | Q92989 | 47615 | | | | 75 | 1314 | 100 |

| | | | Proteolyzed TSEN15–34 | | |
|---|---|---|---|---|---|
| **Protein** | **UniProtKB** | **Mass** (Da) | **Peptide sequences** (#) | **Spectra** (#) | **Sequence coverage** (%) |
| TSEN15 | Q8WW01 | 18629 | 9 | 92 | 87 |
| TSEN34 | Q9BSV6 | 33631 | 14 | 275 | 34 |

**Supplementary Table 3 | Masses of proteolytic fragments of TSEN 15 andTSEN34 obtained from denaturing.** The experimentally determined and theoretically calculated masses as well as the mass difference are given.

| Protein fragment | Experimental mass (Da) | Theoretical mass (Da) | Δ mass (Da) |
|---|---|---|---|
| TSEN15 (residues 23 to 170) | 16313.9±1.0 | 16314.7 | -0.8 |
| TSEN15 (residues 23 to 171) | 16469.8±0.9 | 16470.9 | -1.1 |
| TSEN34 (residues 208 to 310) | 11614.8±0.7 | 11615.2 | -0.4 |

**Supplementary Table 4 | Identification of proteolytic fragments of TSEN15 and TSEN34 by LC-MS/MS.** The position in the protein sequence, the sequence of observed tryptic peptides and the number of spectra for each peptide are given. The preceding and following amino acids are given for each peptide sequence.

| TSEN15 | | |
|---|---|---|
| **Residues** | **Peptide sequence** | **# spectra** |
| 23-45 | R.GFGDGGGAPSWAPEDAWMGTHPK.Y | 22 |
| 46-74 | K.YLEMMELDIGDATQVYVAFLVYLDLMESK.S + Oxidation (M) | 4 |
| 75-118 | K.SWHEVNCVGLPELQLICLVGTEIEGEGLQTVVPTPITASLSHNR.I | 6 |
| 119-127 | R.IREILKASR.K | 7 |
| 121-127 | R.EILKASR.K | 7 |
| 128-154 | R.KLQGDPDLPMSFTLAIVESDSTIVYYK.L | 13 |
| 155-170 | K.LTDGFMLPDPQNISLR.R | 17 |
| 155-171 | K.LTDGFMLPDPQNISLRR.- | 16 |

| TSEN34 | | |
|---|---|---|
| **Residues** | **Peptide sequence** | **# spectra** |
| 204-220 | R.VQSKDWPHAGRPAHELR.Y | 1 |
| 208-220 | K.DWPHAGRPAHELR.Y | 37 |
| 221-230 | R.YSIYRDLWER.G | 61 |
| 231-253 | R.GFFLSAAGKFGGDFLVYPGDPLR.F | 1 |
| 240-253 | K.FGGDFLVYPGDPLR.F | 6 |
| 254-279 | R.FHAHYIAQCWAPEDTIPLQDLVAAGR.L | 7 |
| 280-286 | R.LGTSVRK.T | 4 |
| 280-298 | R.LGTSVRKTLLLCSPQPDGK.V | 8 |
| 286-298 | R.KTLLLCSPQPDGK.V | 75 |
| 286-310 | R.KTLLLCSPQPDGKVVYTSLQWASLQ.- | 1 |
| 287-298 | K.TLLLCSPQPDGK.V | 68 |
| 287-310 | K.TLLLCSPQPDGKVVYTSLQWASLQ.- | 3 |
| 299-310 | K.VVYTSLQWASLQ.- | 3 |

**Supplementary Table 5 | X-ray data collection, refinement, and validation statistics.** The structure of TSEN15–34 was determined from one protein crystal. Values in parentheses are given for highest-resolution shell.

| | TSEN15–34 |
|---|---|
| **Data collection** | |
| Space group | P 1 21 1 |
| Cell dimensions | |
| *a*, *b*, *c* (Å) | 34.85, 69.28, 94.79 |
| $\alpha, \beta, \gamma$ (°) | 90, 98.31, 90 |
| Resolution (Å) | 28.3 - 2.1(2.175 - 2.1) |
| $R_{merge}$ | 0.07428 (0.8863) |
| $I / \sigma I$ | 13.37 (1.44) |
| Completeness (%) | 0.99 (0.99) |
| Redundancy | 5.9 (5.9) |
| **Refinement** | |
| Resolution (Å) | 28.3 - 2.1 |
| No. reflections | 25898 (2576) |
| $R_{work}$ / $R_{free}$ | 19.18 (30.49) / 25.28 (36.27) |
| No. atoms | 3636 |
| Protein | 3516 |
| Ligand/ion | 12 |
| Water | 120 |
| *B*-factor (average, Å$^2$) | 60.17 |
| Protein | 60.10 |
| Ligand/ion | 78.56 |
| Water | 60.32 |
| R.m.s. deviations | |
| Bond lengths (Å) | 0.009 |
| Bond angles (°) | 0.99 |
| **Validation** | |
| Ramachandran plot | |
| Favored (%) | 97 |
| Allowed (%) | 2.5 |
| Outliers (%) | 0.2 |
| Rotamer outliers (%) | 1 |
| Clash score | 9.26 |

**Supplementary Table 6 | DSF data analyzed by ProteoPlex.** $T_d$ – denaturing temperature.

| **TSEN complex** | **$T_d$ - Boltzman (°C)** | **$T_d$ - ProteoPlex (°C)** | **$R^2$ (fit to data)** | **$R^2$ (fit to 2-state unfolding)** |
|---|---|---|---|---|
| TSEN (wt) | 51.0 | 52.1 | 0.99973 | 0.99899 |
| TSEN ($T2^{Y309C}$) | 44.8 | 46.3 | 0.99979 | 0.99955 |
| TSEN ($T34^{R58W}$) | 44.1 | 46.5 | 0.99955 | 0.99902 |
| TSEN ($T54^{S93P}$) | 46.5 | 48.7 | 0.99935 | 0.99818 |
| TSEN ($T54^{A307S}$) | 49.6 | 50.7 | 0.99978 | 0.99951 |

**Supplementary Table 7 | List of patient-derived primary fibroblast cells used in this study.**

| Cell line | Mutations | Description | Zygosity |
|---|---|---|---|
| Ba1 | *TSEN54 c.919G / 919G* | control | homozygous |
| Ba2 | *TSEN54 c.919G / 919G* | control | homozygous |
| Ba3 | *TSEN54 c.919G / 919G* | control | homozygous |
| Ba5 | *TSEN54 c.919G>T / 919G>T* | PCH2 patient | homozygous |
| Ba8 | *TSEN54 c.919G>T / 919G>T* | PCH2 patient | homozygous |
| Ba9 | *TSEN54 c.919G>T / 919G>T* | PCH2 patient | homozygous |
| Ba10 | *TSEN54 c.919G>T / 919G>T* | PCH2 patient | homozygous |
| Ba12 | *TSEN54 c.919G / 919G>T* | parent of Ba19 | heterozygous |
| Ba13 | *TSEN54 c.919G / 919G* | control | homozygous |
| Ba14 | *TSEN54 c.919G / 919G* | control | homozygous |
| Ba15 | *TSEN54 c.919G / 919G* | control | homozygous |
| Ba17 | *TSEN54 c.919G / 919G>T* | parent of Ba19 | heterozygous |
| Ba18 | *TSEN54 c.919G>T / 919G>T* | PCH2 patient | homozygous |
| Ba19 | *TSEN54 c.919G>T / 919G>T* | PCH2 patient | homozygous |
| Ba20 | *TSEN54 c.919G>T / 923delC* p.(Pro318Gln fsX23) | PCH4 patient | compound heterozygous |
| Ba245 | *TSEN54 c.919G>T / 919G>T* | PCH2 patient | homozygous |
| Ba1230 | *TSEN54 c.919G>T / 919G>T* | PCH2 patient | homozygous |
| Ba1613 | *TSEN54 c.919G / 919G>T* | parent of Ba1597 | heterozygous |
| Ba1614 | *TSEN54 c.919G / 919G>T* | parent of Ba1597 | heterozygous |
| Ba1597 | *TSEN54 c.919G>T / 919G>T* | PCH2 patient | homozygous |
| T1 (BAB3846) | *CLP1 c.419G / 419G>A* | parent of BAB3402 | heterozygous |
| T3 (BAB3402) | *CLP1 c.419G>A / 419G>A* | patient | homozygous |

**Supplementary Table 8. Hydro-tRNAseq data (separate file).**

### Source Data

**Source Data 1 | Uncropped images as shown in Fig. 1.**

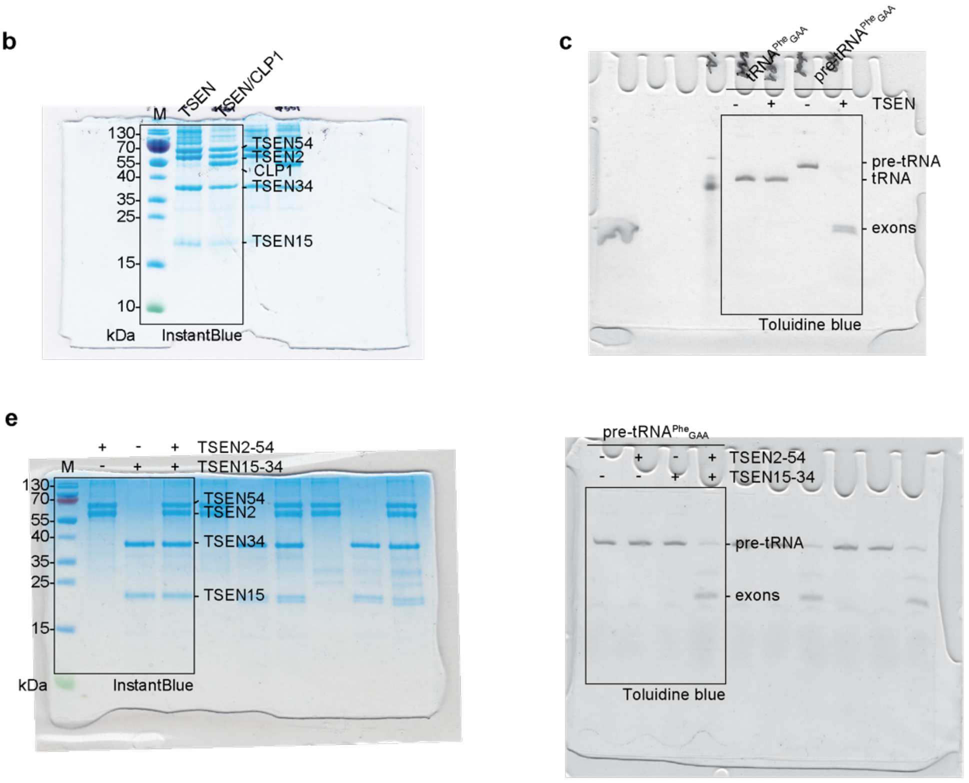

**Source Data 2 | Uncropped images as shown in Extended Data Fig. 1.**

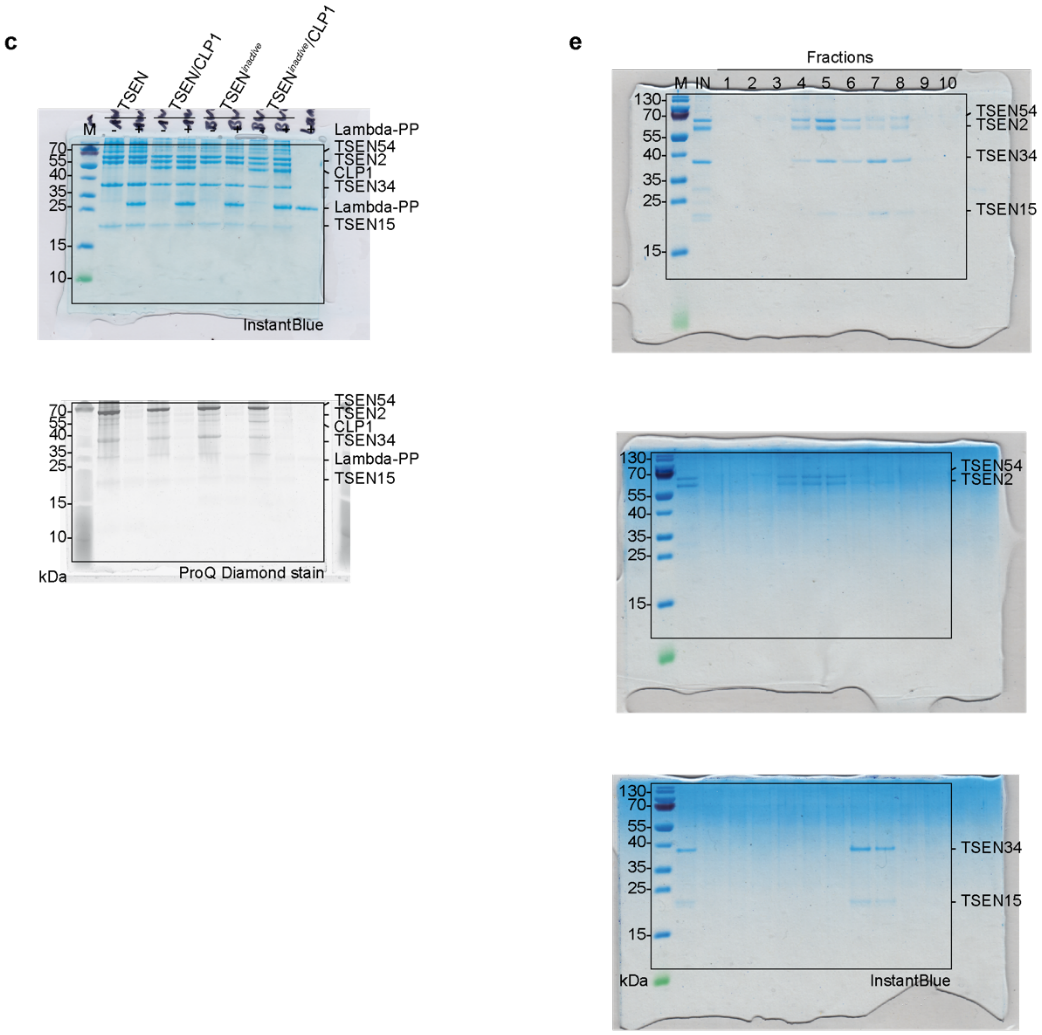

**Source Data 3 | Uncropped images as shown in Fig. 2.**

a

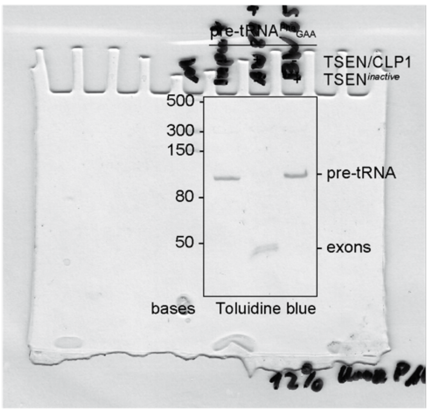

b

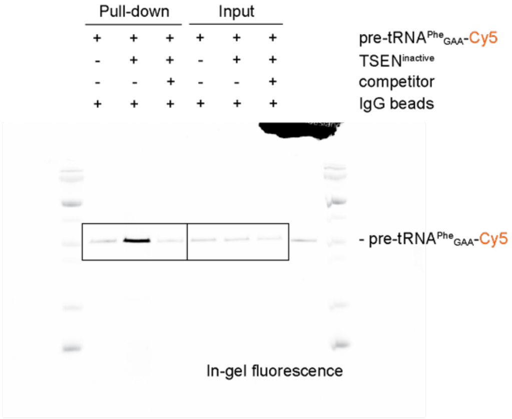

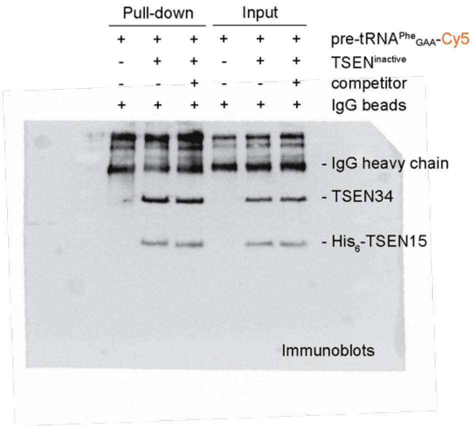

f

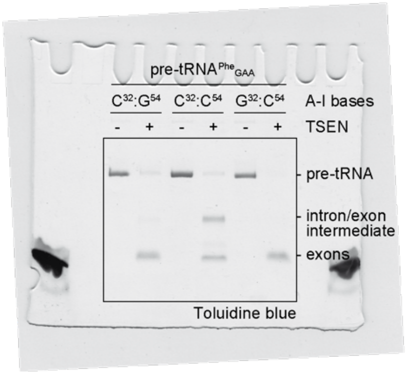

**Source Data 4 | Uncropped images as shown in Extended Data Fig. 2.**

a

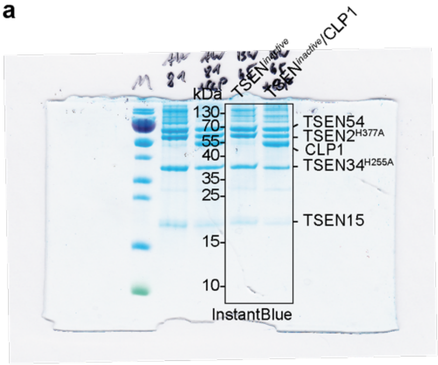

b

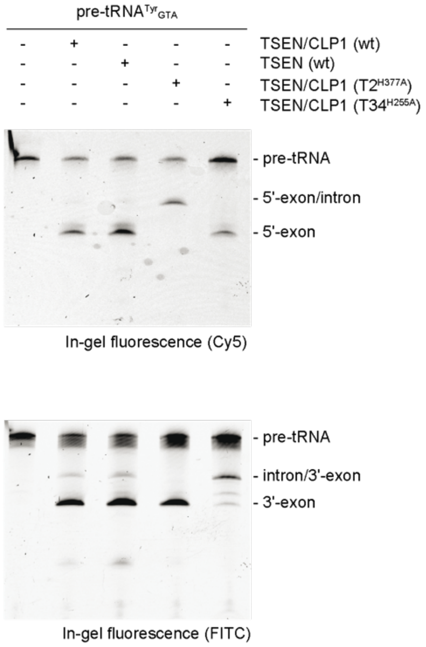

d

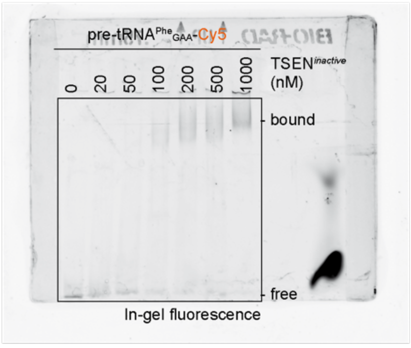

e

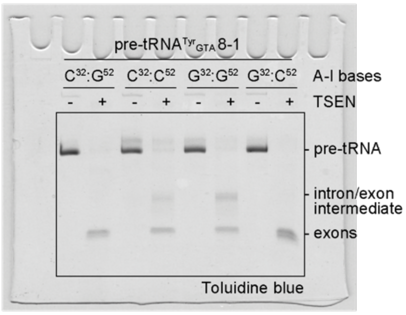

**Source Data 5 | Uncropped images as shown in Fig. 3.**

f

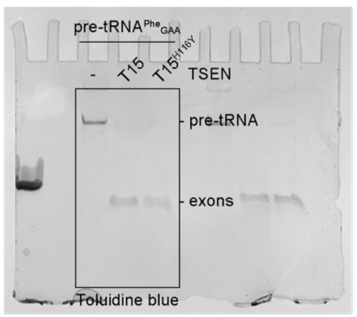

e

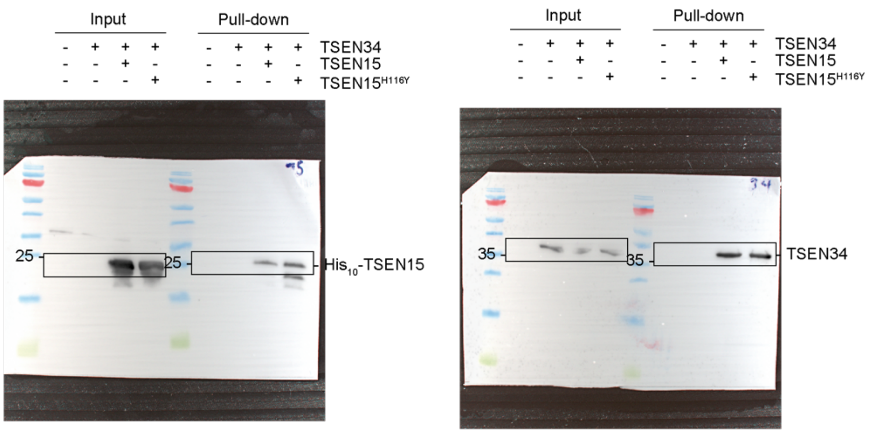

**Source Data 6 | Uncropped images as shown in Extended Data Fig. 3.**

**a**

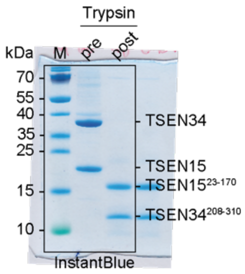

**d**

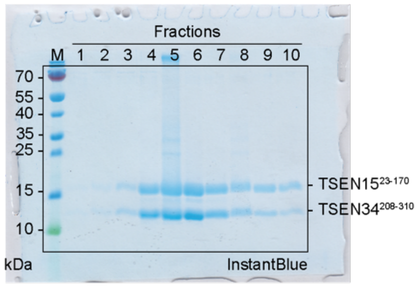

**Source Data 7 | Uncropped images as shown in Fig. 4a.**

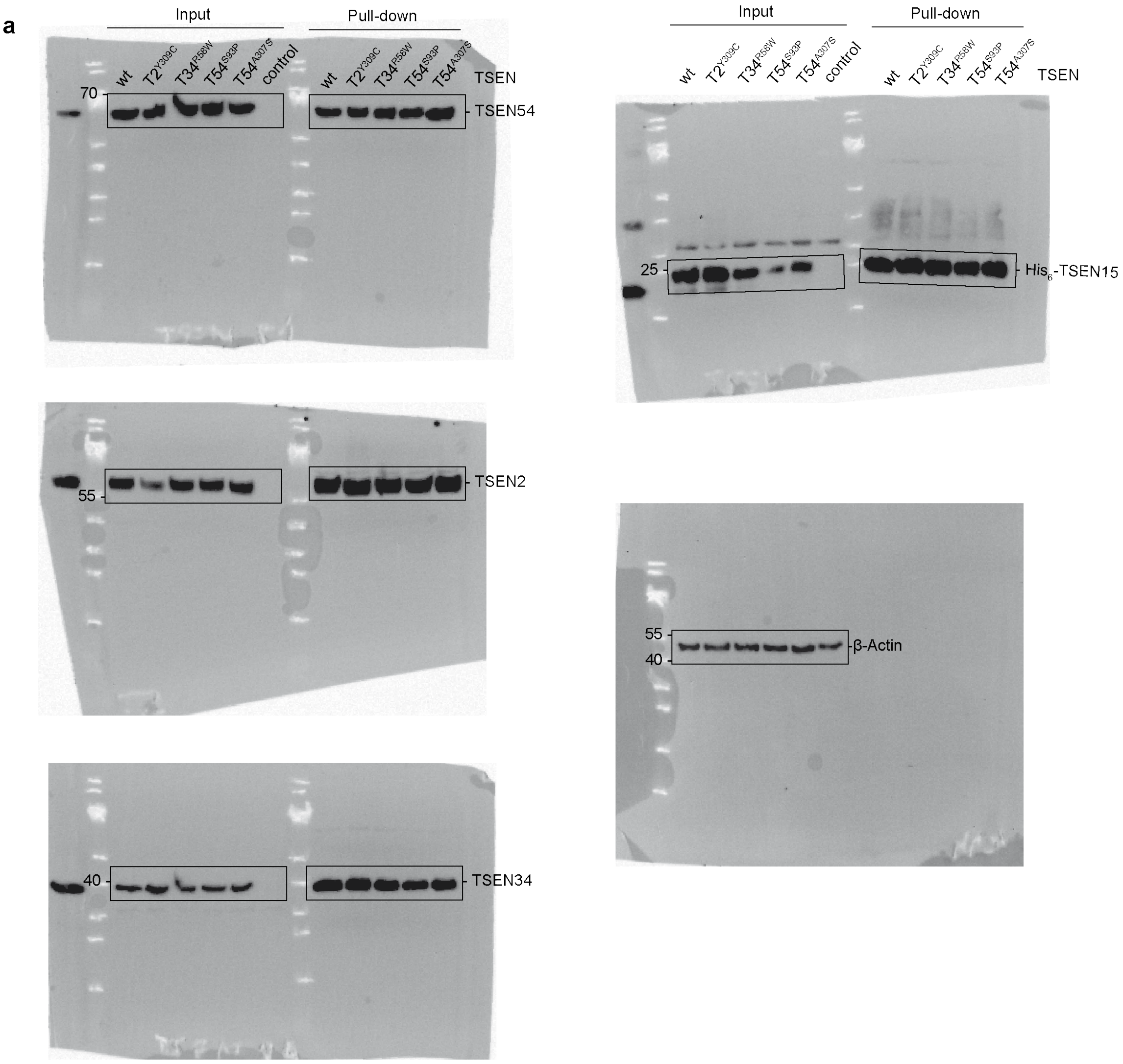

**Source Data 8 | Uncropped images as shown in Fig. 4b,c.**

**b**

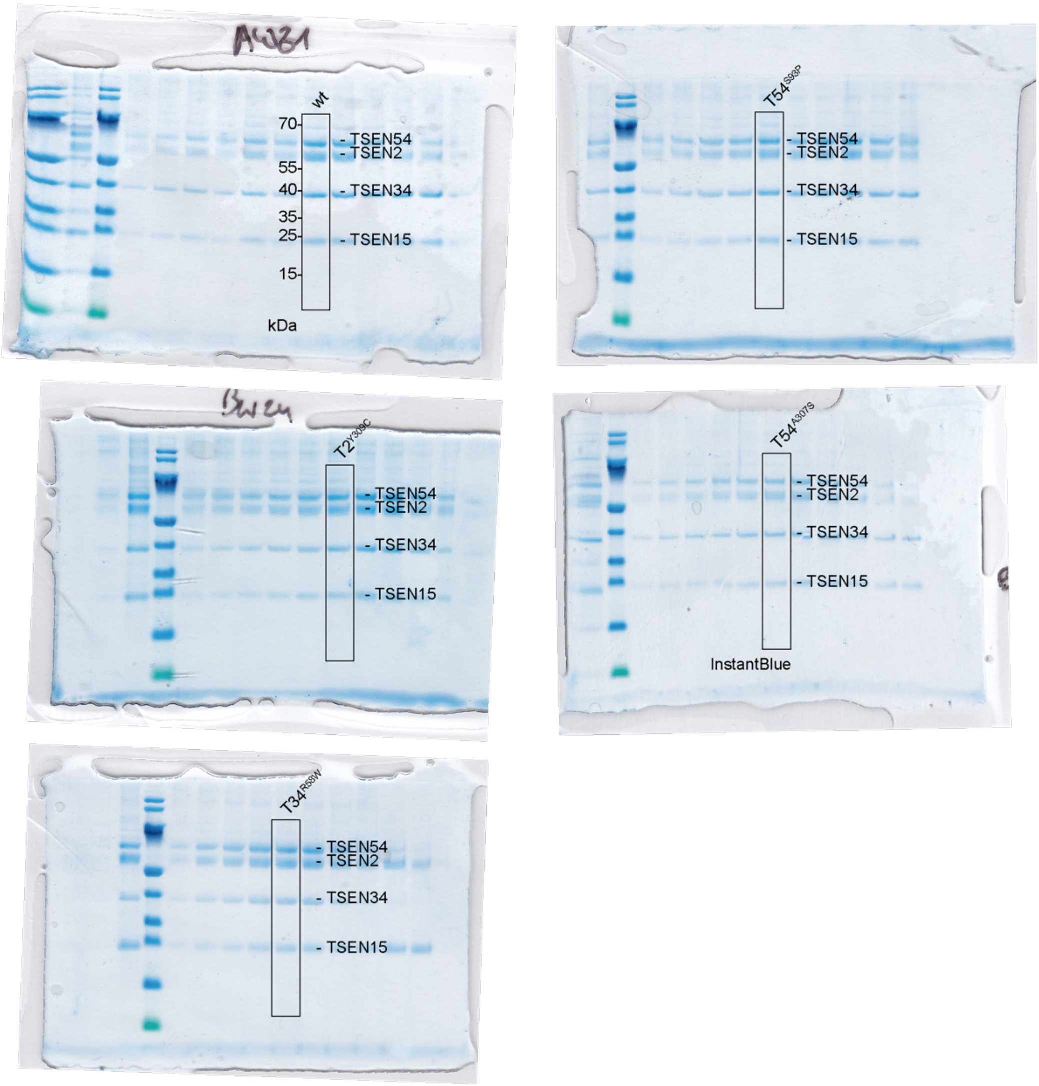

**c**

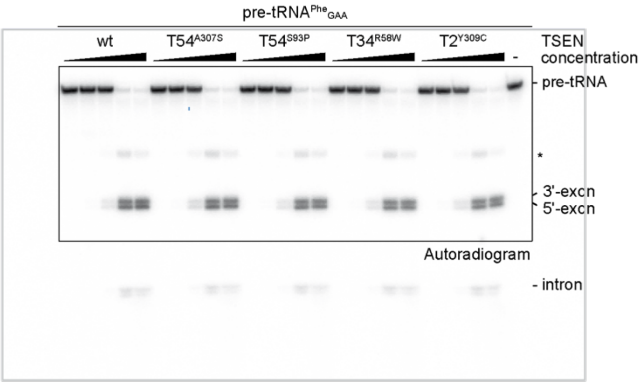

**Source Data 9 | Uncropped images as shown in Extended Data Fig. 5.**

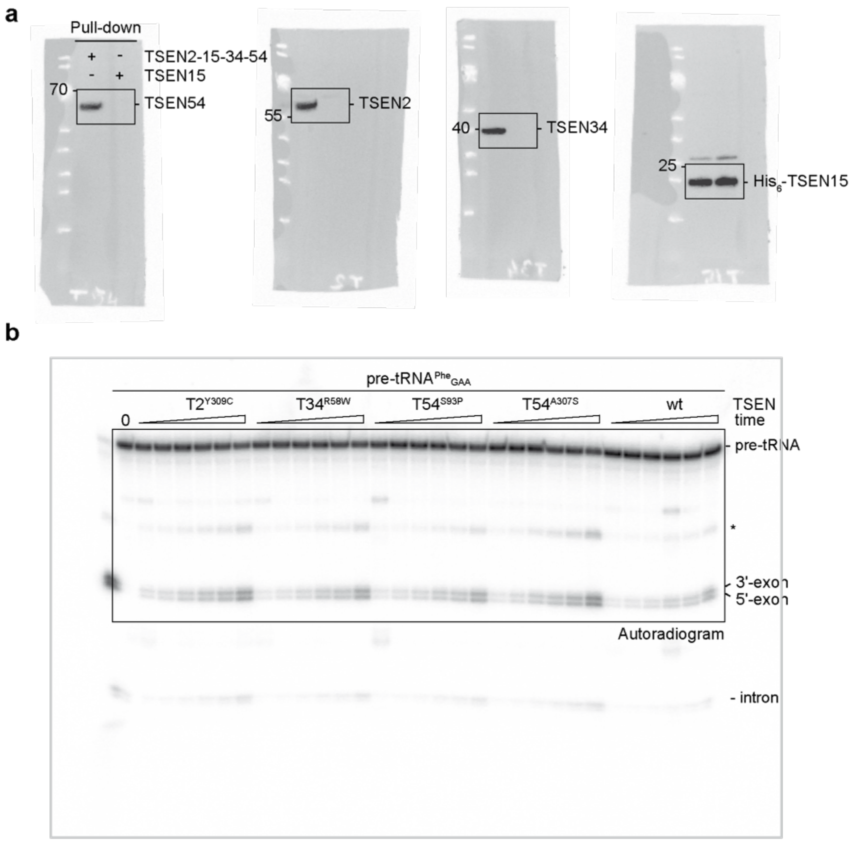

**Source Data 10 | Uncropped images as shown in Fig. 5.**

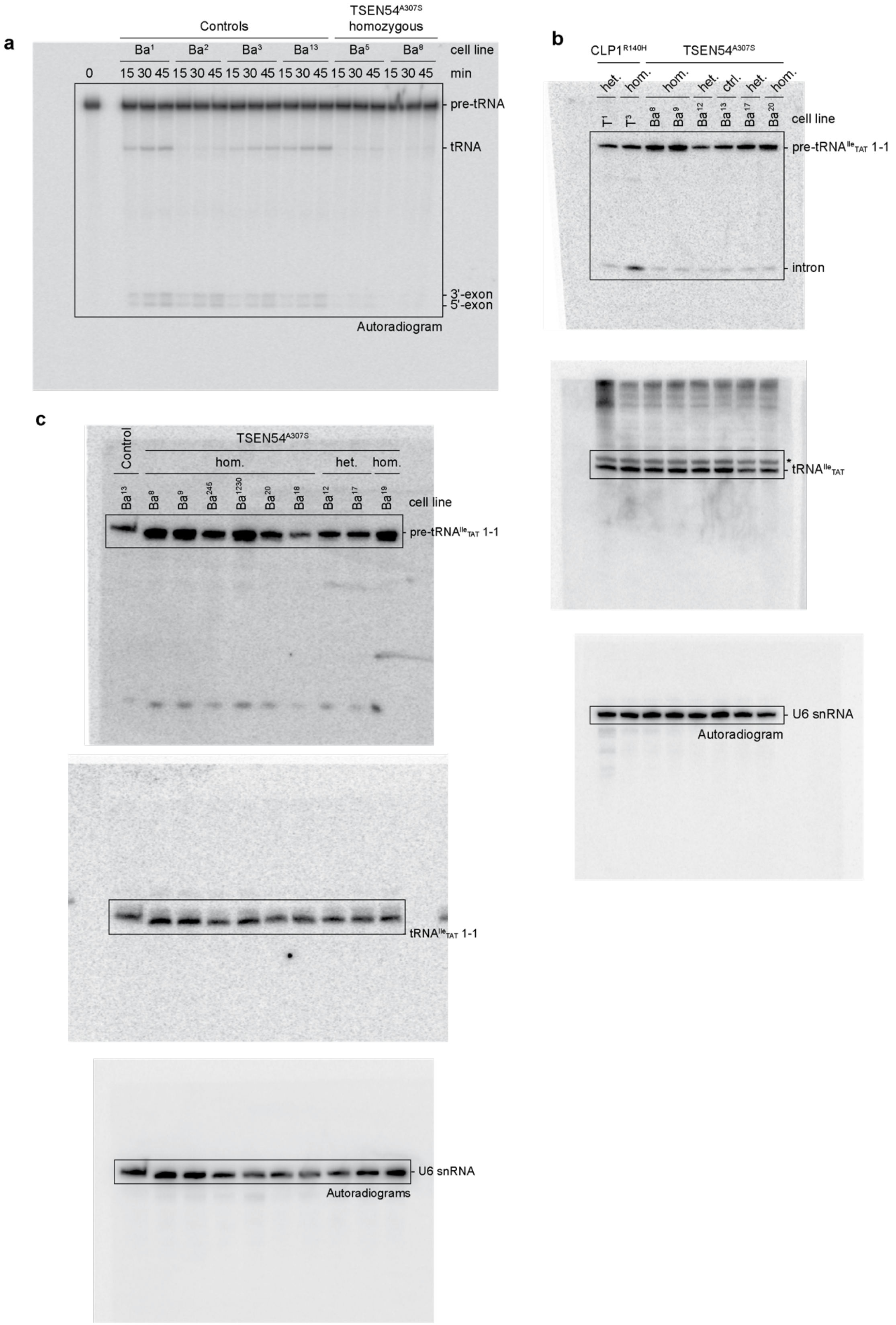

**Source Data 11 | Uncropped images as shown in Fig. 6a.**

**a**

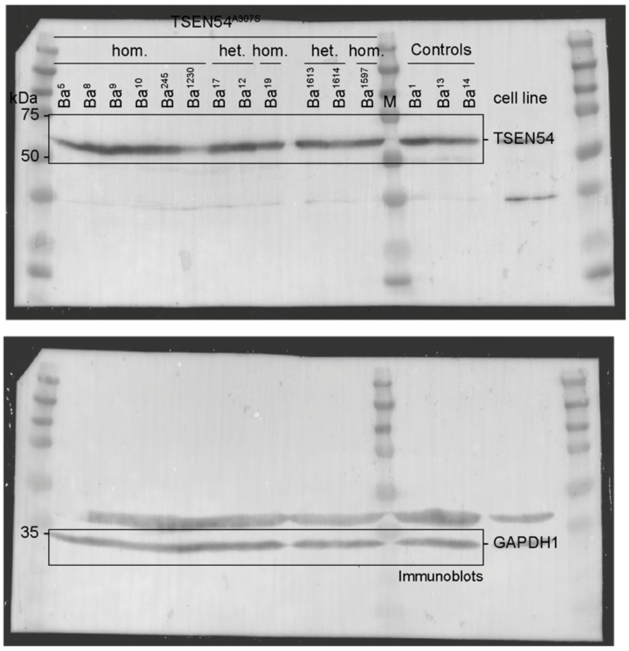

105 **Source Data 12 | Uncropped images as shown in Fig. 6b.**

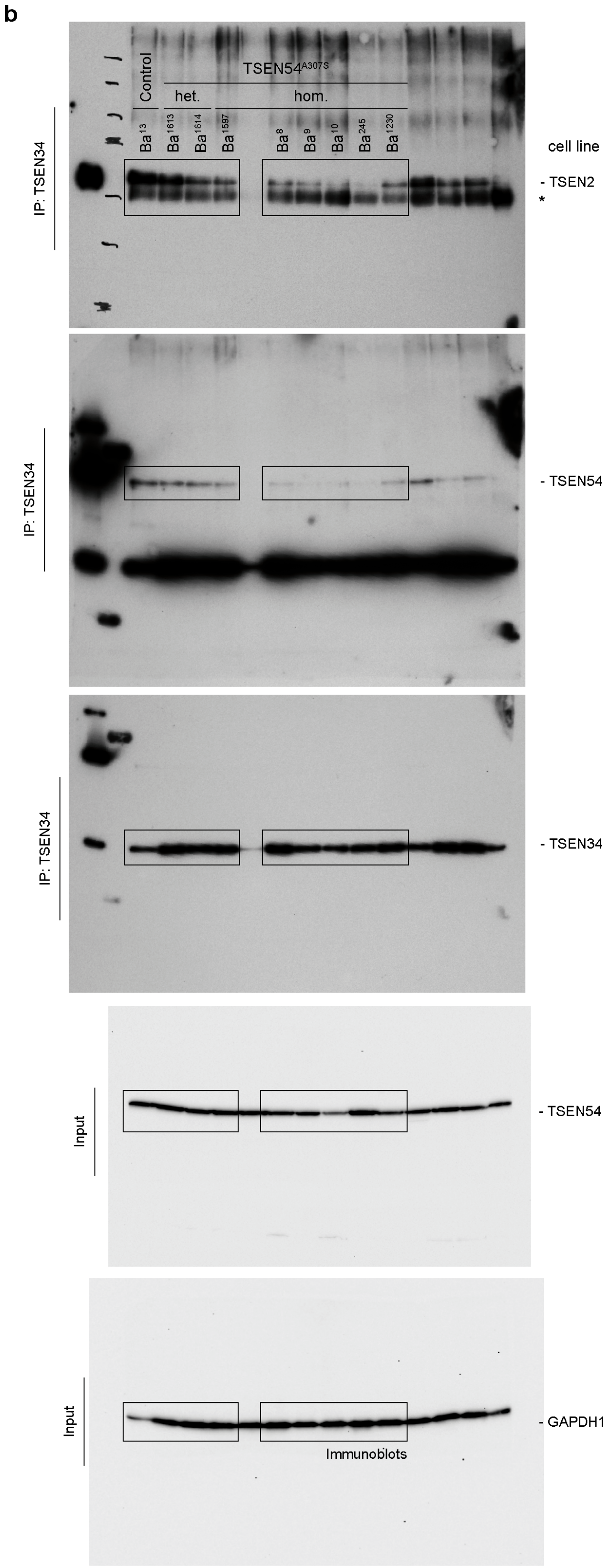

106
107

**Source Data 13 | Uncropped images as shown in Fig. 6c.**

c

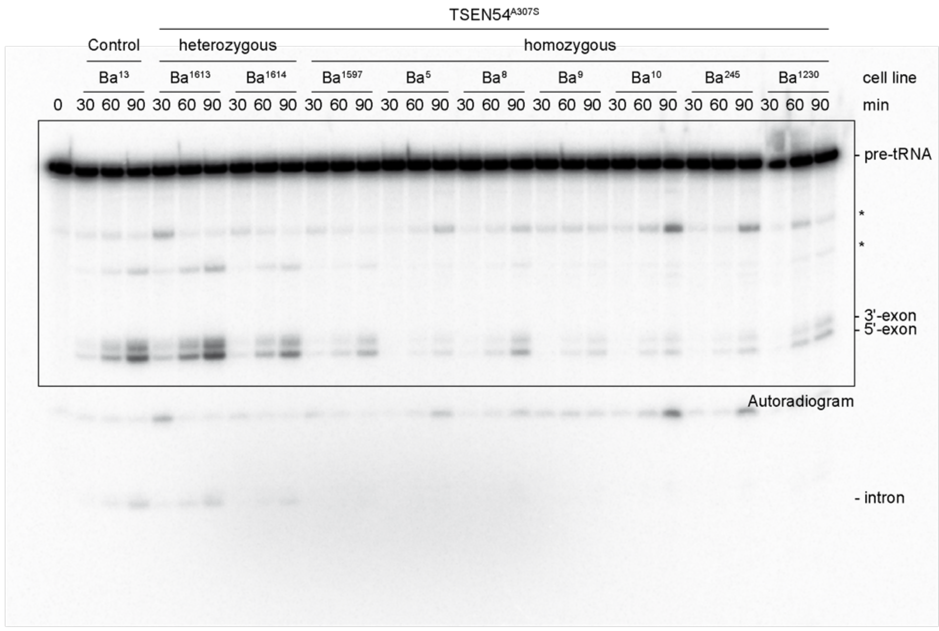

*Gamma-corrected image*

111 **Source Data 14 | Uncropped images as shown in Extended Data Fig. 6a.**

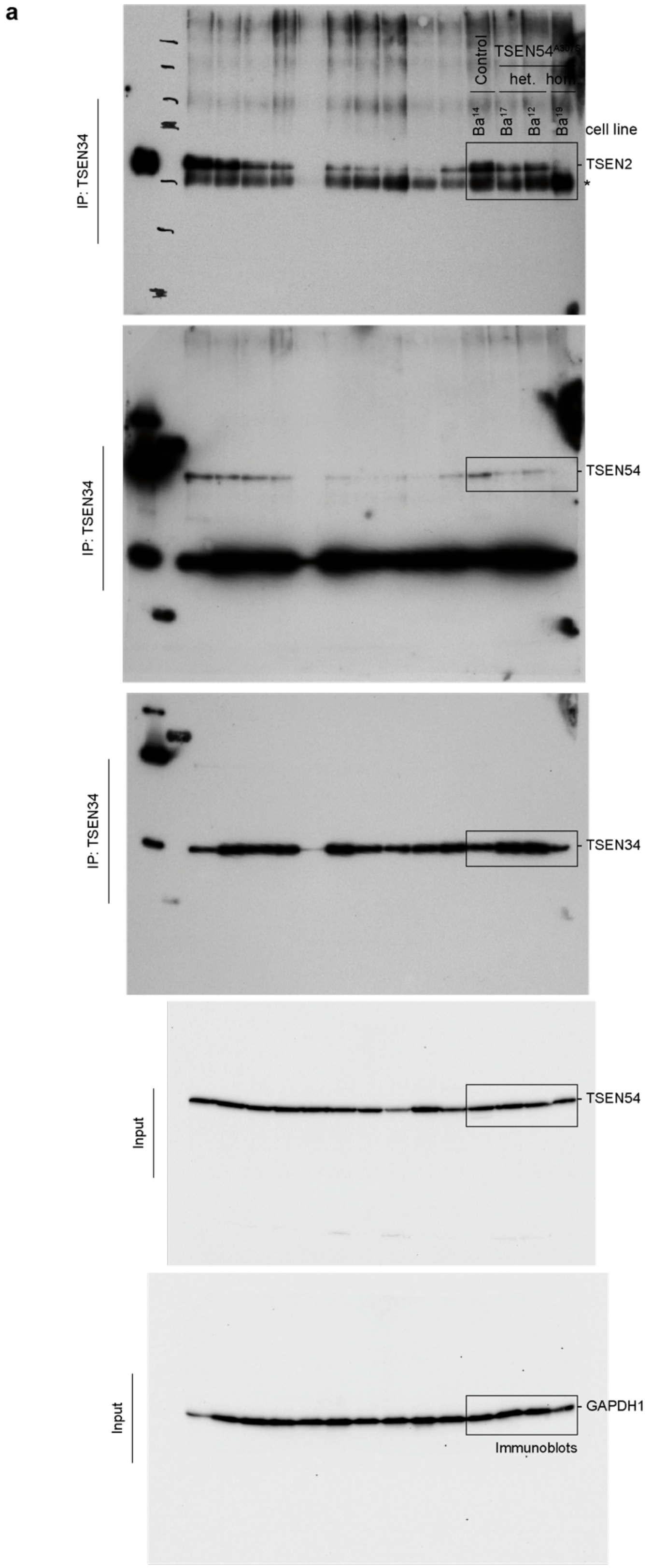

112
113

**Source Data 15 | Uncropped images as shown in Extended Data Fig. 6b,c.**

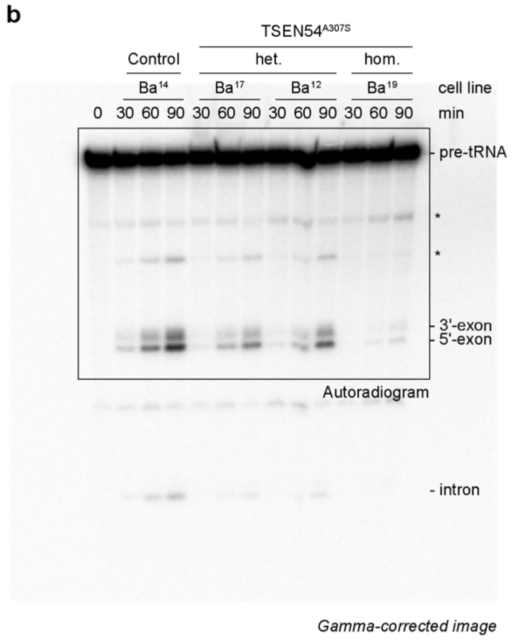

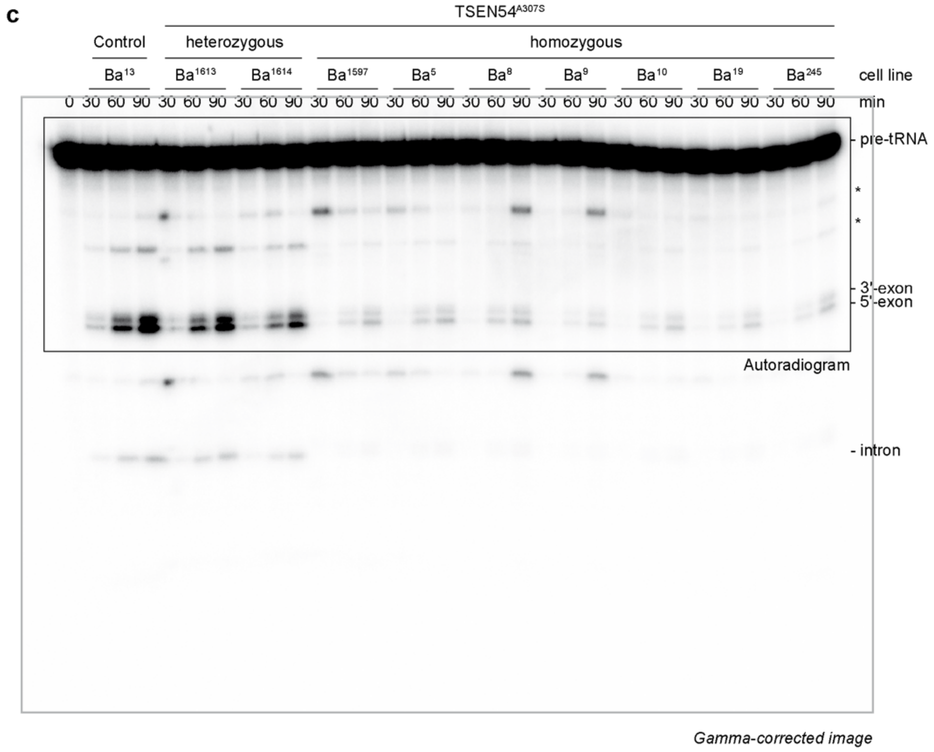
